# End-to-end evaluation of pipelines for metagenome-assembled genomes reveals hidden performance gaps

**DOI:** 10.64898/2026.04.06.712906

**Authors:** Izaak Coleman, Jiong Ma, Gordon Qian, Yusheng Jiang, Aya Brown Kav, Tal Korem

## Abstract

The generation of Metagenome Assembled Genomes (MAGs) has become a standard and basic step in the analysis of metagenomic data. This multi-step process, which includes assembly, binning, refinement, and quality control, has many alternative approaches, algorithms, and parameters. Determining the ideal approach for a given ecosystem and study, or highlighting algorithmic gaps in need of additional research and development, requires rigorous benchmarking. We present MAG-E (**MAG** pipeline **E**valuator), a generalizable and expandable framework for end-to-end evaluation of entire MAG pipelines: from assembly, through binning, to quality control and filtering. MAG-E relies on simulations that are built to match an ecosystem of interest and provide a ground truth for accurate evaluation. To demonstrate the capabilities of MAG-E, we benchmark two assemblers, six binning algorithms, three binning modes, and three quality control and refinement methods in the context of the human gut microbiome. Our findings offer multiple insights into optimal MAG generation in this context. We find that metaSPAdes consistently outperforms MEGAHIT in terms of recall (completeness), and that COMEBin overall outperforms alternative binning algorithms, but has lower precision than SemiBin2. While multi-sample binning results in higher precision, as previously shown, single-sample binning has higher recall and leads to better overall performance with modern binners. Binning refinement, which combines bins from multiple different algorithms, leads to reduced performance. We further show that CheckM2 systematically overestimates completeness and underestimates contamination, and that this is partially ameliorated when using GUNC. Finally, we analyze performance at the contig level, and demonstrate that binning algorithms systematically underperform for prophages and fail to bin contigs that are shared between genomes. Overall, MAG-E offers deep insights into successes and gaps in this important analytic process.

## Introduction

Metagenomic sequencing enables the study of genomes in their native environment, bypassing the need for isolation and culturing. However, current sequencing technologies and assembly algorithms cannot yet produce complete genomes, instead providing shorter genomic fragments (“contigs”). To better recover the microbial genomes from which these contigs originated, it is common practice to cluster contigs based on sequence composition and coverage, using algorithms colloquially referred to as “binners”. The “bins” these algorithms create are then often refined and quality-filtered, producing Metagenome-Assembled Genomes (MAGs). This process has become a fundamental step in metagenomic data analysis, instrumental to comprehensively charting ecosystems such as the vaginal and gut microbiomes^1–3^, analyzing pangenomes at scale^4^, characterizing new branches of the tree of life^5,6^, and training genome language models^7^. Many different binning algorithms have been proposed^8–13^, each with different parameters and settings; they have been applied either to each sample separately^14–16^ or to multiple samples at a time^2,17^; and different approaches have been used to refine and quality check the resulting bins^1,2,15^. This constantly expanding space creates a need for robust evaluation approaches that enable investigators to choose the best methods and parameters for analyzing their datasets.

Many approaches have so far been taken to evaluate the MAG generation process, with three key properties differentiating between them: (a) <u>Simulated vs. real:</u> whether the metagenomic data used for evaluation is simulated or derived from a genuine microbiome sample; (b) <u>Ground-truth vs. heuristic</u> <u>evaluation:</u> whether evaluation is based on exact knowledge of each sample’s content, or whether it is based on estimates, such as those produced by CheckM2^18^; and (c) <u>Sample complexity:</u> whether the evaluation reflects the complexity and abundance distribution of the microbial community that is of interest. Each of these properties has a critical impact on evaluation: simulated samples provide access to ground-truth information, while such ground-truth is available for complex “real” samples only with intensive efforts (e.g., extensive isolate sequencing^19^ or Hi-C^20^); heuristic evaluation has been shown to underestimate contamination^17^; and sample complexity has been shown to impact binning performance^21^. Other important considerations include which steps of the MAG generation process are evaluated and whether contig-specific biases are studied. For example, while many studies evaluated different binning algorithms^19,21–24^, only one compared different quality-control methods^17^, and none evaluated the impact of assembly on binning. Previous evaluations have found that contig properties, such as GC content or the presence of repetitive elements, affect the accuracy of binning^19,24^, highlighting existing gaps in the field. A best-effort summary of past benchmarking efforts in light of these properties is provided in **Supplementary Table 1.**

To address these considerations, we present MAG-E (**MAG** pipeline **E**valuator), a generalizable framework for ecosystem-specific evaluation of MAG generation pipelines. MAG-E constructs realistic simulations that match the complexity of an ecosystem of interest at the strain level while providing an accurate ground truth that facilitates a comprehensive evaluation based on isolate genomes. Simulating gut metagenomic samples, we evaluate different steps of the MAG generation pipeline, including assembly, binning, refinement, and quality control. We further examine contig-specific properties to identify systematic biases in binning performance. MAG-E is expandable to additional methods and applicable to multiple ecosystems, thereby providing an easy-to-use framework for developers to benchmark their tools and for investigators to select the best tool for their ecosystem of interest.

## Results

### Constructing realistic and complex simulated metagenomes

To conduct a ground-truth-based evaluation of MAG construction pipelines, we developed a metagenomic simulator that simulates realistic samples matching the complexity of an ecosystem of interest. MAG-E therefore takes as input a real metagenomic sample and, using a highly accurate metagenome profiler^25^, creates a ground-truth specification that “mirrors” it using isolate genomes and MAG sequences from a provided database. We select references to match the species-level composition of the input sample (at 95% ANI), as well as subspecies strain diversity (at 98% ANI) when it is detected (e.g., “Species A” in **Fig. 1a**; **Methods**). Because isolate genomes are more complete^19^ and harder to bin (two-sided Wald z-test *p*<10^−6^; **Methods**; **Fig. 1b**; **Table S2**), we preferentially select them over MAGs for inclusion in simulations when multiple options exist for a strain (**Fig. 1a**). After creating a ground-truth specification, we use InSilicoSeq^26^ to simulate the sample at the same sequencing depth, sampling reads from the genomes in the ground-truth specification according to their inferred abundance levels of the input sample. MAG-E can then run multiple MAG pipelines, including different assemblers, binning algorithms in different modes (single-sample, multi-sample, etc.), binning refiners, and quality-control methods, and then evaluate their outputs against the ground truth. We assign recall, precision, and F-score for each ground-truth genome in every sample (**Fig. 1c**; **Methods**). While reference MAGs are included in the simulated sample to increase its complexity, our evaluation uses only ground-truth isolate genomes. MAG-E provides a simple API that facilitates the use of different assemblers and binners and their application to different ecosystems and databases (**Methods**).

**Figure 1.**
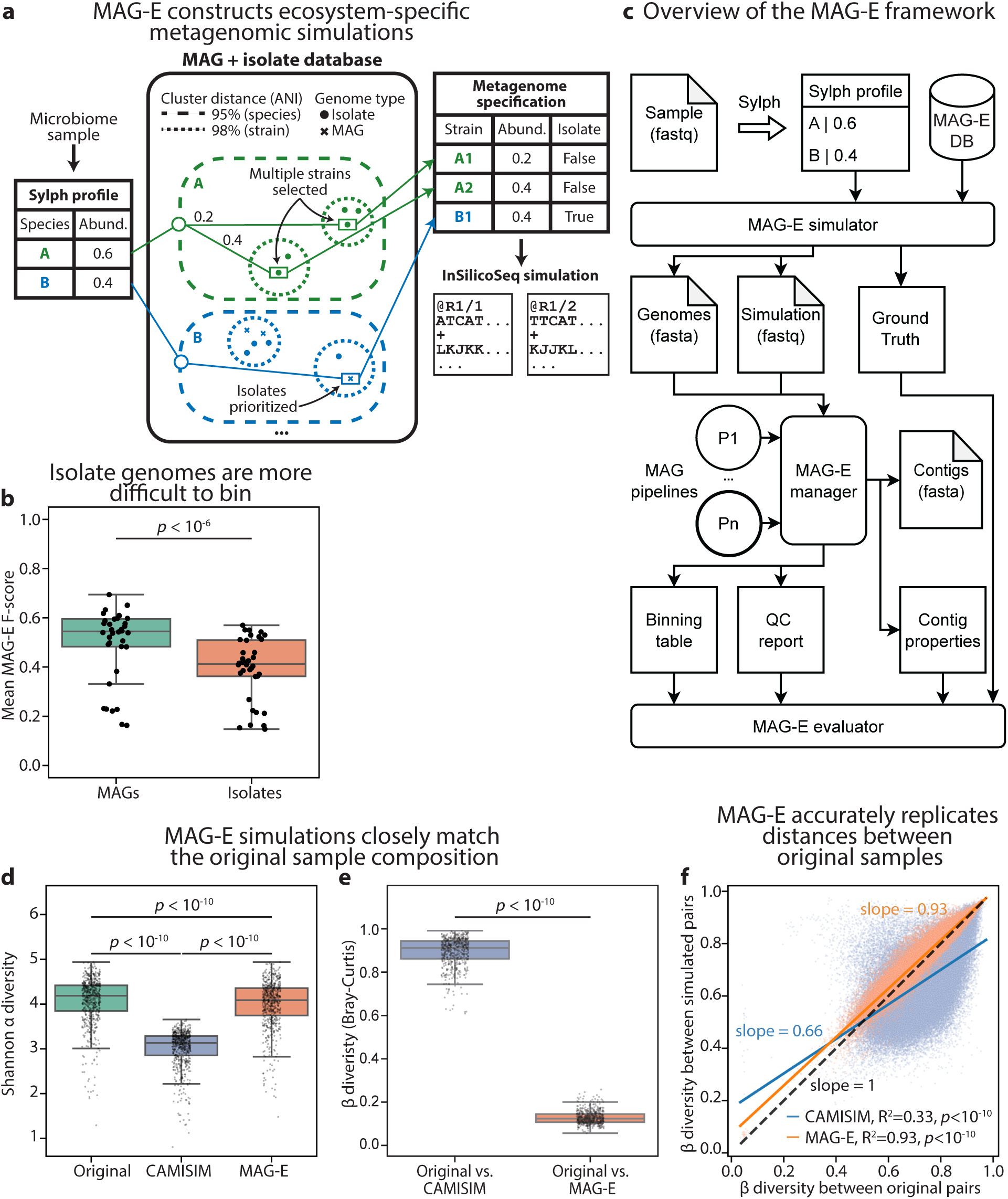
MAG-E simulates realistic metagenomic data with complexity matching input samples. **a,** An illustration of the MAG-E simulation process. For a given microbiome sample, we use Sylph^25^ to infer the species profile with respect to a database containing MAGs and isolate genomes (e.g., UHGG^1^). We then run an additional Sylph query against all the genomes within each species and select the strain cluster (98% ANI) with the best match as a representative for the species, allowing the selection of additional strain clusters in case of additional high-confidence matches. From each such strain cluster, we select a reference genome, preferring isolates over MAGs and otherwise prioritizing ANI. We then simulate a metagenome using InSilicoSeq^26^, ensuring that species abundances and read depth match the original sample. See **Methods** for full details. **b,** Mean F-score of 36 MAG pipelines (2 assemblers x 6 binners, excluding DAS Tool x 3 binning modes) over 100 simulated gut metagenomes (**Methods**), comparing between evaluation based on MAGs and isolate reference genomes. **c,** An overview of MAG-E: the MAG-E simulator provides a simulated metagenome and ground-truth genomes for each sample. We then run multiple MAG pipelines for each sample, each including assembly, binning, refinement, and quality control. We use BLAST^27^ to map each ground-truth genome to its best-matching (highest recall) bin, allowing us to calculate evaluation metrics. See **Methods** for full details. **d,** Shannon α diversity for 575 human gut metagenomes, calculated using mOTUs^28^, compared to the α diversity of the corresponding simulations. **e,** Bray-Curtis β diversity between samples simulated by CAMISIM and MAG-E and the corresponding original sample for each. **f,** The β diversity of all pairs of simulated gut metagenomes (y-axis) plotted vs. the β diversity of the corresponding pair of original gut metagenomes (x-axis), shown separately for MAG-E (orange) and CAMISIM (blue). The line of best fit is drawn for each method. Box, IQR; line, median; whiskers, 1.5xIQR; *p*, two-sided Wald z-test (**Methods**; c), Wilcoxon signed-rank test (d,e) or Student’s t-test (f).

Because the gut microbiome is an extensively studied and complex ecosystem, we demonstrate the use of MAG-E using a healthy gut metagenomic cohort^29^ along with the Unified Human Gastrointestinal Genome (UHGG) database^1^ (**Methods**). To evaluate our simulations, we compared them to CAMISIM^30^. We created simulated metagenomes for 575 gut samples^29^ using both CAMISIM and MAG-E, and compared each simulation to its original sample, using taxonomic profiles obtained from mOTUs^28^ (**Methods**). While both methods simulated samples with lower α diversity than the original ones (Wilcoxon signed-rank *p*<10^−10^ for both; **Fig. 1d**), MAG-E-simulated samples had higher α diversity than those simulated by CAMISIM (*p*<10^−10^; **Fig. 1d**), and smaller absolute differences from the α diversity of the original samples (*p*<10^−10^; **Fig. S1a**). Similarly, the β diversity between the composition of the original and MAG-E simulated samples was significantly lower than between the original and CAMISIM-simulated samples (*p*<10^−10^; **Fig. 1e**). Finally, to assess how well the simulations capture the overall population structure, we evaluated all sample pairs, and compared the β diversity between pairs of original samples to the β diversity between the corresponding simulated pairs. We found that MAG-E captured this population-level β diversity structure very accurately (R^2^=0.93; slope of 0.93; Student’s t-test *p*<10^−10^), while CAMISIM led to a substantial underestimation of β diversity (R^2^=0.33; slope of 0.66; *p*<10^−10^; **Fig. 1f**). Similar results were obtained when we compared reference-free Mash distances^31^ rather than β diversity (**Fig. S1b,c**). Overall, these results demonstrate that MAG-E produces highly realistic simulations that closely match input samples.

### COMEBin, SemiBin2 and metaSPAdes outperform alternatives for MAG generation

To comprehensively evaluate different steps of the MAG generation process, we evaluated 36 pipelines using a subset of 100 samples: two assemblers, MEGAHIT^32^ and metaSPAdes^33^; six binners, CONCOCT^10^, MaxBin2^9^, METABAT2^8^, VAMB^11^, SemiBin2^13^, and COMEBin^12^; and three binning modes: single-sample, multi-sample^17^, and a partial multi-sample approach in which a subset of similar samples is used^2^; for a total of 3,600 MAG-generation tasks. We selected a set of 808 ground-truth genomes that were recovered from their simulated samples, defined as ≥0.7 recall and ≥0.9 precision by at least one MAG pipeline. We then evaluated all MAG pipelines on a per-genome basis using recall (completeness), 1-precision (contamination), and F-score (**Fig. 2a,b; Fig. S2a; Fig. S3**), and used a linear mixed model that accounts for the structure of the data to evaluate performance (**Methods; Tables S3-6**). Comparing the performance of the two assemblers we evaluated, we found that metaSPAdes led to higher recall and F-score (two-sided Wald z-test *p*<10^−6^ for both; **Figs. 2c, S2b**). This may be related to the larger size of metasSPAdes assemblies (mean±std of 2.5×10^8^±0.76×10^8^ and 2.1×10^8^±0.68×10^8^ basepairs for metaSPAdes and MEGAHIT; Wilcoxon signed-rank *p*<10^−10^; **Fig. S4a**) despite their smaller N50 (mean±std of 2702±1700 and 3951±2448 basepairs; *p*<10^−10^; **Fig. S4b**). Both assemblers had similar levels of precision (Wald z-test *p*=0.19; **Fig. 2d**), which is expected, as contamination is primarily the result of misbinning and not of assembly quality.

**Figure 2.**
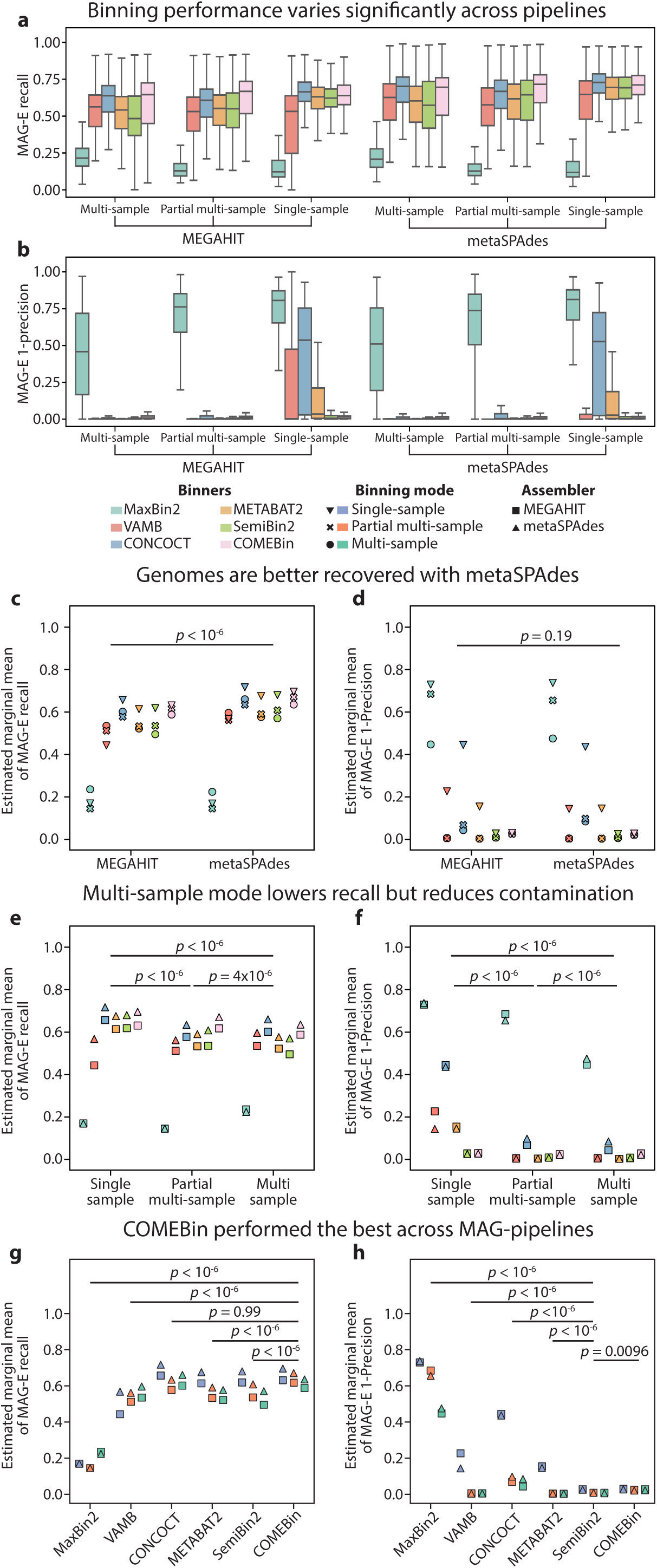
Evaluation of MAG pipeline performance reveals tradeoffs between recall and precision. **a-f,** Results pertain to analysis of 36 MAG pipelines (two assemblers, six binners and three binning modes) on a set of 808 recoverable genomes (**Methods**). **a,b**, Boxplots (box, IQR; line, median; whiskers, 1.5xIQR) showing MAG-E recall (a) and 1-precision (b) of the recoverable genomes across all samples for each pipeline. **c-h**, Estimated marginal mean MAG-E recall (c,e,g) and 1-precision (d,f,h), stratified by assembler (c,d), binning mode (e,f), and binner (g,h), obtained by fitting linear mixed models to results at the genome level (**Methods**; **Tables S3-5**), and *p*-values are Tukey HSD adjusted two-sided Wald z-tests of differences in estimated marginal means for the relevant comparisons (**Methods**).

Recently, binning mode has been associated with contamination, with a claim that multi-sample binning is superior to single-sample binning as it reduces contamination by using co-abundance across samples to further differentiate between contigs with similar coverage^17^. Examining binning modes (**Fig. 2e,f**), we observed the same, with single-sample binning leading to higher contamination (1-precision; two-sided Wald z-test *p*<10^−6^, Tukey HSD adjusted; **Fig. 2f**) and lower F-scores (*p*<10^−6^; **Fig. S2c**). However, this was not consistent across all binners, with SemiBin2 and COMEBin showing minimal difference between binning modes (**Fig. 2f**). Furthermore, lower contamination for multi-sample binning came at the cost of lower recall (*p*<10^−6^ for single-sample vs. either partial multi-sample or multi-sample **Fig. 2e**). Overall, our results suggest that in scenarios in which recovering genomes to their fullest extent is more important than the precision of each bin, investigators could consider using single-sample binning.

Finally, we evaluated the performance of every binning algorithm. We found that MaxBin2 underperformed all other binners, in terms of recall, precision, and F-score (**Figs. 2g,h, S2d**). CONCOCT and COMEBin had significantly higher recall than other binners (Tukey HSD adjusted two-sided Wald z-test *p*<10^−6^ for either COMEBin or CONCOCT vs. all others; *p*=0.99, for comparison between them; **Fig. 2g**), and the best two pipelines in terms of recall were metaSPAdes assembly followed by binning with either single-sample CONCOCT or single-sample COMEBin (estimated marginal mean MAG-E recall of 0.72 and 0.70, respectively; *p*<0.05 for either vs. all other pipelines except two for COMEBin; *p*=0.28 for comparison between the two; **Tables S3,S4**). However, CONCOCT had very high contamination when run in single-sample mode following either MEGAHIT or metaSPAdes assembly (marginal precision of 0.56 for both; **Fig. 2h; Tables S3,S5**). SemiBin2 had the best precision (lowest contamination; *p*=0.0096 vs. COMEBin, *p*<10^−6^ vs. all others; **Fig. 2h**). Summarizing over recall and precision using F-scores, COMEBin was significantly better than other binners (*p*<10^−6^ for all; **Fig. S2d**). The best pipelines of the 48 we tested used metaSPAdes assembly followed by single-sample COMEBin, single-sample SemiBin2, or partial multi-sample COMEBin (estimated marginal mean F-score of 0.81, 0.80 and 0.79, respectively; *p*<0.05 for either vs. all other pipelines; p>0.6 between these top three; **Fig. S2**; **Tables S3,6**).

### Binning refinement using DAS Tool led to reduced performance

Binning refiners are often used to combine the results of different binning algorithms^34^. To explore their performance, we evaluated DAS Tool^34^, a popular binning refiner, using two approaches: “Nayfach”, which followed Nayfach et al. ^15^ and integrated binning solutions from MaxBin2, METABAT2, and CONCOCT; and “Best”, which integrated binning solutions from the three binners with highest F-score: METABAT2, SemiBin2, and COMEBin (**Fig. S2d; Table S6**). We evaluated 12 DAS pipelines, with each using binners that were run following the same assembler and in the same binning mode, for a total of 1,200 MAG-generation tasks. While DAS(Nayfach) improved upon results obtained using MaxBin2 (Tukey HSD adjusted two-sided Wald z-test *p*<10^−6^ for recall, precision, or F-score), it obtained worse results than METABAT2 and CONCOCT (p<10^−6^ for both, comparing either recall, precision, or F-score; **Figs. 3a,b, S5, S6a**). Similarly, DAS(Best) obtained significantly worse results than each of the individual binners it operated on (p<10^−6^ for all comparisons; **Figs. 3c,d, S5, S6b**). Overall, our results demonstrate that binning refinement does not necessarily improve binning.

**Figure 3.**
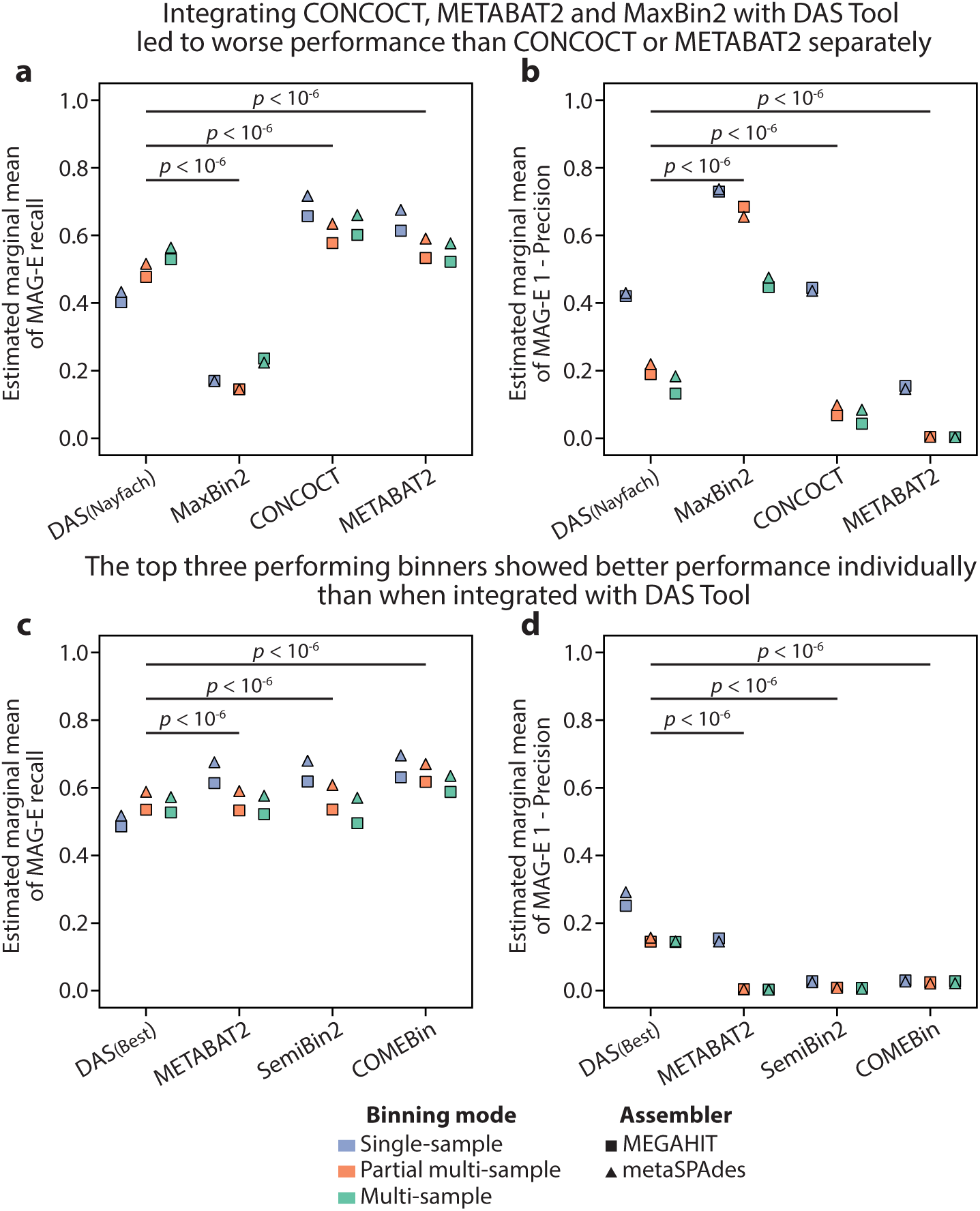
Individual binners outperformed DAS Tool. **a,b,** Estimated marginal mean MAG-E recall (a) and 1-precision (b) for DAS(Nayfach) pipelines, which combine the binning outputs of MaxBin2, CONCOCT, and METABAT2 pipelines (also shown). **c,d,** same as (a,b) but for DAS(Best) pipelines, which combine the outputs of METABAT2, SemiBin2, and COMEBin. Estimated marginal means obtained by fitting linear mixed models to results at the genome level (**Methods**; **Tables S3-5**), and *p*-values are Tukey HSD adjusted two-sided Wald z-tests of differences in estimated marginal means for the relevant comparisons (**Methods**).

### Contig-level analysis reveals systematic loss of shared genomic elements

Recent studies revealed that binning algorithms often mis-bin repeated, mobile, or short contigs^19,20,24^. MAG-E provides ground-truth assignments at the contig level, facilitating an analysis of factors underlying lack of binning. To demonstrate this, we analyzed all contigs from our recoverable genomes set, and examined contig-level properties affecting their recall (**Methods**), To avoid genome length bias when looking at subsets of contigs, we calculated “assembly recall”^35^, which considers only the assembled portion of a genome, rather than its entire sequence. Considering METABAT2, COMEBin and SemiBin2 in multi-sample mode following assembly with metaSPAdes, we found that contigs whose coverage and sequence composition (tetra-nucleotide compositions) differed from the mean coverage and sequence composition of their originating genomes had lower recall (**Fig. 4a,b**), which is expected given that these are the main properties used by binners. We note that all three binners had higher tolerance for difference in sequence composition than in coverage, with contigs at the third quartile of sequence composition similarity still having similar recall to those in the first quartile (**Fig. 4b**). Mobile genetic elements often have outlying coverage and sequence composition. Examining such contigs specifically, we found systematic lower recall for contigs annotated as prophages by geNomad^36^, as well as for shared contigs that mapped to more than one ground-truth genomes (Tukey HSD adjusted two-sided Wald z-test *p*<10^−6^ for all comparisons; **Fig. 4c; Table S7**). Consistent with our genome-level analysis, COMEBin had better recall for both contig groups (*p*<10^−6^ for all comparisons).

**Figure 4.**
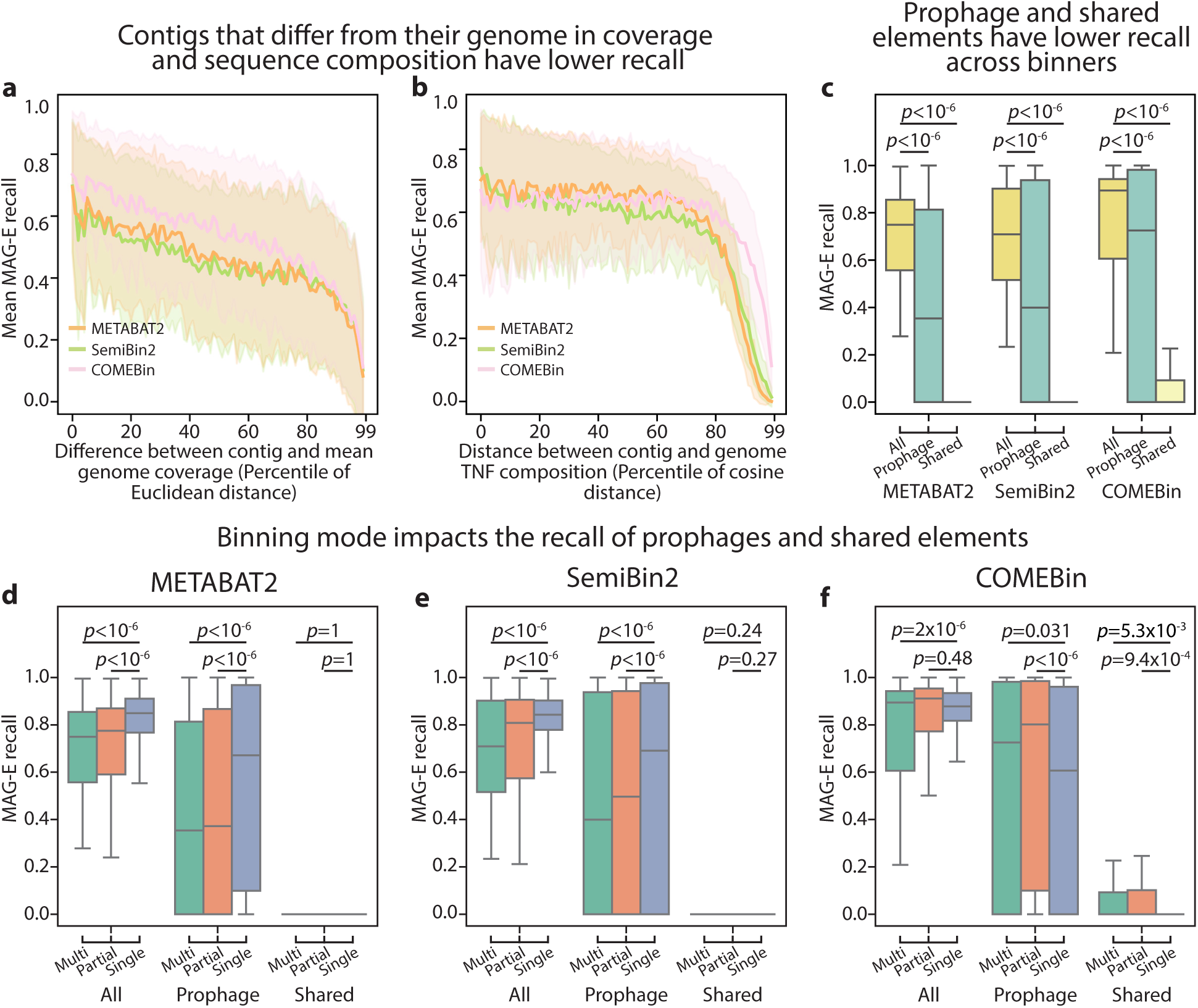
Contig-level analysis reveals binner-specific interaction between binning mode and recovery of variable genomic regions. Results pertain to analysis of contig-level recall of MAG pipelines using metaSPAdes followed by COMEBin^12^, METABAT2^8^, and SemiBin2^13^, ran in multi-sample mode (a-c) or all three modes (d-f). **a,** MAG-E mean recall (y-axis) over percentile bins calculated by comparing the Euclidean distance between the coverage of each contig and the mean coverage of all contigs assigned to its ground-truth genome across all samples. **b,** Same as (a), but for the cosine distance between the tetranucleotide (TN) composition of the contig and that of its ground-truth genome. **c,** MAG-E recall of each recoverable genome in each sample, stratified by contig type and binner. **d-f,** MAG-E recall of each recoverable genome in each sample, stratified by binning mode and contig type, for METABAT2 (d), SemiBin2 (e) and COMEBin (f). a,b: line, mean; shaded region, 1 s.d. c-f: box, IQR; line, median; whiskers, 1.5xIQR. *p*, Tukey HSD adjusted two-sided Wald z-test.

Multi-sample binning observes co-abundance of genetic elements across samples, and therefore might miss variable genomic regions. We therefore next compared between the three binning modes (single-sample, multi-sample, and partial multi-sample) for the same three binners (METABAT2, COMEBin, and SemiBin2). Interestingly, while METABAT2 and SemiBin2 indeed showed higher recall for prophages in single-sample binning (Tukey HSD adjusted two-sided Wald z-tests *p*<10^−6^, for both binners in single-sample vs. other modes; **Fig. 4d,e; Table S8,9**), COMEBin had lower recall for these elements using single-sample binning (*p*=0.031 and *p*<10^−6^ for single-sample vs. multi-sample and partial multi-sample respectively). All three tools recovered very few shared contigs in either mode, but COMEBin did have significantly higher recovery in both multi-sample and partial multi-sample mode compared to single-sample (*p*=0.005 and 9×10^−3^, respectively; **Fig. 4d-f; Table S8-10**). Such insights may guide both investigators who are pursuing specific genomic analyses and algorithm developers who wish to identify bottlenecks for further improvements.

### Quality control methods overestimate completeness and underestimate contamination

It is an accepted standard in the field^37^ to classify bins as low, medium, and high quality (LQ, MQ and HQ, respectively) according to estimates of completeness (<50%, ≥50%, and >90%) and contamination (<10%, <10% and <5%). To examine the performance of quality control, we ran CheckM2^18^ on bins with ≥2.5x coverage (**Methods**) produced by the different pipelines, disregarding MaxBin2 due to poor performance and lack of HQ bins in two of the pipelines, for an overall of 64,596 bins from 3,000 MAG-generation tasks. Using our ground truth to evaluate the estimated marginal mean of MAG-E recall, 1-precision, and F-score for genomes whose bins fell into each of the categories for each of the 30 MAG pipelines, we found that these classifications indeed separated bins by quality: HQ bins had higher recall than MQ (Tukey HSD adjusted two-sided Wald z-test *p*<10^−6^**, Methods**), and MQ higher than LQ (*p*<10^−6^; **Fig. 5a**). Similarly, HQ bins had lower 1-precision than MQ (*p*<10^−6^), and MQ lower than LQ (*p*<10^−6^; **Fig. 5b**). However, CheckM2 overestimated the completeness of MQ and HQ bins (estimated marginal mean recall of 0.45 and 0.62, respectively), with none of the MAG pipelines obtaining marginal mean recall over 0.9 for HQ bins (**Fig. 5a**). Furthermore, 18 and all 30 of the MAG pipelines yielded a marginal mean 1-precision over 10% and 5% for MQ and HQ bins, respectively (estimated marginal mean 1-precision of 0.13 and 0.10; **Fig. 5a,b; Table S11**). Examining one pipeline more closely (metaSPAdes followed by multi-sample COMEBin) we found that while CheckM2 consistently over-estimated recall, most outliers had low precision (high contamination; **Fig.5c**; **Table S12**). Similarly, while CheckM2 underestimated contamination for a substantial number of bins, it tended to do so for those with lower recall (**Fig. 5d**). Examining these results across multiple MAG pipelines demonstrated similar general trends, along with pipeline-specific patterns (**Figs. S7-12**), likely driven by the quality of the bins generated by each pipeline. Our results concur with previous analyses^17,38^ that demonstrated that quality assessment algorithms may systematically underestimate contamination.

**Figure 5.**
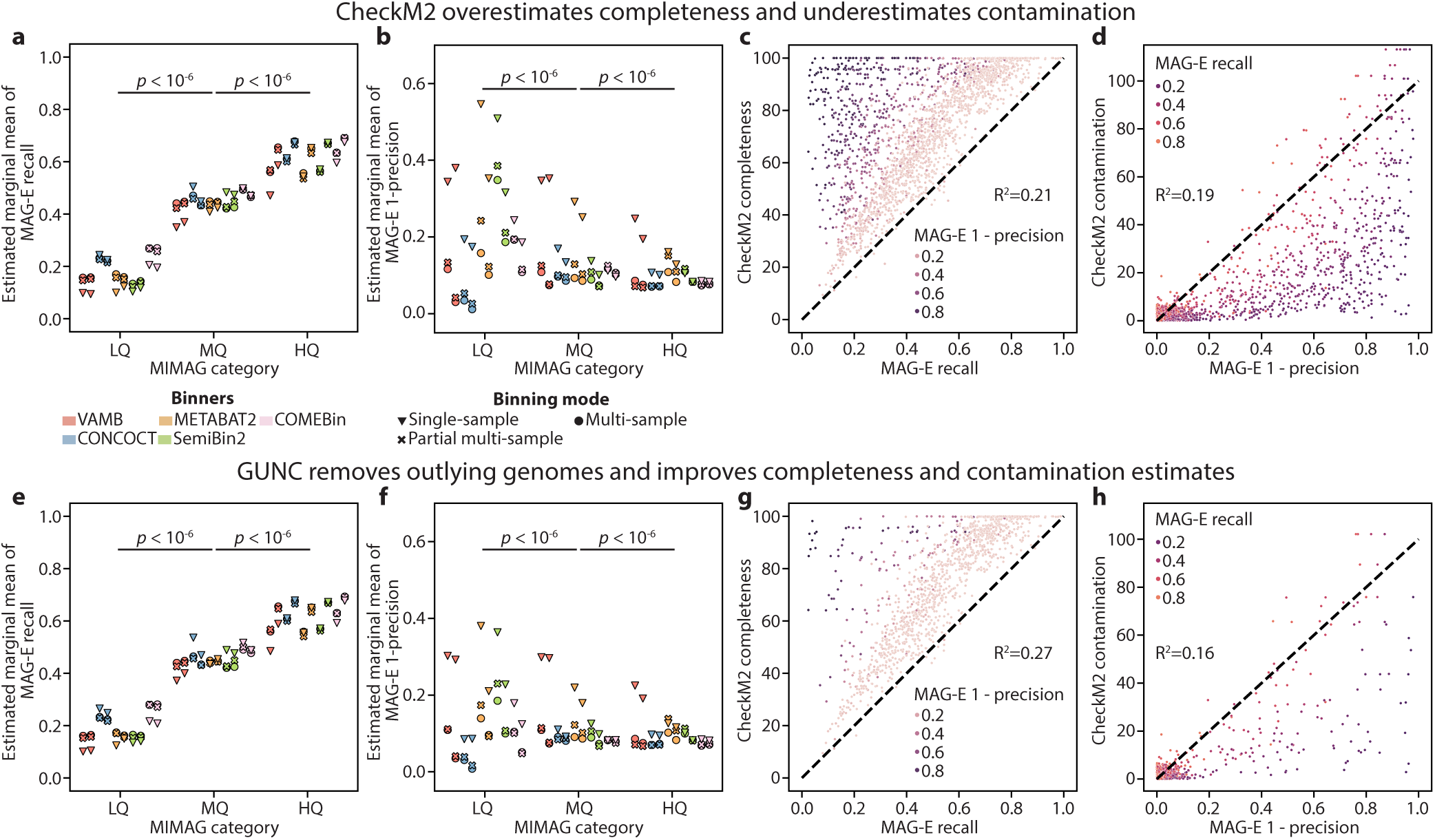
CheckM2 overestimates recall and precision. **a,b,** Estimated marginal mean MAG-E recall (a) and 1-precision (b) for each MAG pipeline, stratified by classification^37^ to low (LQ; <50% completeness and <10% contamination), medium (MQ; ≥50% and <10%) and high quality (HQ; >90% and <5%) MIMAG categories by CheckM2^18^. **c,d,** CheckM2-estimated completeness (y-axis, c) and contamination (y-axis, d) vs. MAG-E recall (c) and 1-precision (d) for every ground-truth genome (N=2,284) processed by the MAG pipeline that included metaSPAdes and COMEBin in multi-sample mode (**Methods**). Shown is the marginal coefficient of determination (R^2^) from a linear mixed model with a random effect for sample and genome. Note that CheckM2 contamination estimates can exceed 100%. **e-h,** Same as a-d, respectively, after removal of bins flagged by GUNC^38^ as contaminated. Box, IQR; line, median; whiskers, 1.5xIQR; *p*, two-sided Wald z-test, Tukey HSD adjusted.

Several tools have been developed which evaluate contamination via approaches that are complementary to CheckM^15,38–40^. Prominent of these is GUNC^38^, which predicts the taxonomic lineage of each gene in a bin, uses inconsistent lineage assignments between contigs as indication of contamination, and is often used in conjunction with CheckM2^2,41–43^. We therefore next evaluated whether filtering bins with GUNC improves binning quality. GUNC identified a mean±std of 584±404 bins per pipeline as contaminated (26.5%±17.3% of bins), which had a mean±std MAG-E recall of 0.28±0.25 and 1-precision of 0.47±0.61. Interestingly, its application improved CheckM2 estimates: it improved the recall of bins classified as LQ and MQ (Tukey HSD adjusted two-sided Wald z-test p<10^−6^ for comparing pre- and post-GUNC for both; **Fig. 5e**), as well as their precision (p<10^−6^; **Fig. 5f; Table S11**). However, the same systematic overestimation of completeness was still observed for MQ and HQ bins (estimated marginal mean recall of 0.45 and 0.62, respectively; **Fig. 5e**). Examining the same specific MAG pipeline as before (in **Fig. 5c,d**), we found that GUNC filtered many of the bins for which CheckM2 overestimated completeness and underestimated recall, but this did not lead to substantial changes to CheckM2’s accuracy (R^2^ increased from 0.21 to 0.27 for completeness and decreased from 0.19 to 0.16 for contamination; **Fig. 5g,h; Table S12**). Similar trends were observed across most pipelines (**Fig. S7-12; Table S12**).

## Discussion

The variety of available assembly and binning algorithms, each with different parameters and running modes, and paired with various choices for refinement and quality control, join together combinatorially to create a vast space of analytic options. Investigators therefore need a reliable way to benchmark the best choice for their study, and algorithm developers need ways to highlight current limitations and bottlenecks as well as to benchmark new tools. To address this, we developed MAG-E, a framework for benchmarking the full gamut of the MAG generation pipeline. MAG-E relies on simulations that are realistic, match a specific ecosystem of interest, and provide a ground truth for accurate evaluation. We demonstrated an application of MAG-E for evaluating MAG pipelines for the gut metagenome. We found several surprising insights, including that CheckM2, a widely used tool, overestimated completeness and underestimated contamination; that the metaSPAdes assembler had superior recall to MEGAHIT despite MEGAHIT’s higher N50; that the aggregation of different binning results using DAS Tools led to a decrease in performance; and that while in general multi-sample binning outperformed single-sample, the best pipeline combinations used COMEBin and SemiBin2 in single-sample mode. Evaluating binning performance at the contig level, we found a systematic bias against contigs that are shared across genomes or that contain prophage elements, demonstrating a clear and substantial methodological gap in the field.

MAG-E simulates metagenomic samples with a ground truth, which are meant to match the complexity of a particular sample. An alternative approach is to run binners or MAG pipelines on real samples and use quality-control tools such as CheckM^18^ to evaluate binning quality^17,44^ (**Table S1**). Our results, which show underestimation of contamination and overestimation of completeness by CheckM2, emphasize the importance of using accurate ground truth information, also highlighted by others^35^. Nonetheless, simulation approaches are limited by the accuracy and realism of the simulations themselves. We showed that MAG-E generates better simulations than CAMISIM^30^, the current state of the art in the field. While CAMISIM uses a similar conceptual approach to MAG-E for simulating metagenomes^30^, there are a few notable differences: (a) CAMISIM only uses complete genomes, while MAG-E focuses on isolate genomes, and uses MAGs only when no isolate is available; (b) CAMISIM may include genomes from a different species that share a higher taxonomic rank (by default, up to the family level), while MAG-E will not; and (c) CAMISIM randomly selects the number of strain genomes to use, while MAG-E uses evidence provided from a Sylph^25^ query. Beyond the realism of simulations, MAG-E also allows the user to create a custom database, ensuring its suitability to an ecosystem of interest and otherwise allowing continuous updates to benchmarking efforts. These differences likely explain the improved simulation accuracy by MAG-E.

MAG-E is designed to evaluate MAG generation pipelines in the context of the multiple steps that they involve, rather than focusing just on binning. Our analysis thus includes the first systematic evaluation of CheckM2 and the impact of combining it with GUNC, an evaluation of binning refinement, and a systematic evaluation of contig-level biases. While past studies have demonstrated biases against mobile genetic elements^24^, repetitive sequences^19^, and outlying GC content^24^, this was not done in the context of large-scale simulations with systematic evaluation across multiple tools. We showed that these comparisons are valuable, in that different tools operated better for prophage elements under different binning modes. Our results further highlight a substantial gap in the field in terms of the ability of different binners to handle contigs from the accessory genomes - those that are variable or shared across different genomes. These are particularly important for comparative genomics and population genetics.

Our evaluation approach has a few salient differences from methods applied previously^35,45^. BinBencher supports binning evaluation at higher taxonomic ranks than the genome level, arguing that contamination between highly similar bins could be discounted in certain scenarios^35^. While we agree with these arguments, we have chosen to evaluate MAG generation pipelines at their highest resolution, as improvements in such tasks are likely to also improve evaluation at higher taxonomic ranks. We note that MAG-E addresses other issues highlighted by Nissen et al.^35^, such as allowing contigs to map to any number of ground-truth genomes, correctly accounting for the same genome when it appears in more than one sample, and correctly evaluating multi-sample binning. AMBER^45^ differs from MAG-E in how it calculates performance metrics. AMBER calculates average purity and completeness over the bins produced for a given sample rather than the genomes it contains. Since both the genomes binned and number of bins can vary between binners, MAG-E uses only the ground-truth per-genome performance metrics when evaluating performance. Furthermore, while AMBER provides an evaluation of binning performance using per-sample averages, MAG-E evaluates pipeline performance using per-dataset averages which it calculates using statistical models that analyze per-genome performance while accounting for the dataset’s structure.

An important consideration for future development of MAG-E and other benchmarking frameworks is data leakage. For example, CheckM2^18^ uses a machine learning architecture trained on genomes from RefSeq^46^, which is highly informative of the genomes included in UHGG^1^. We therefore consider our benchmark of CheckM2 to be an upper bound on its performance on completely unseen taxa. The two best performing binners in our analysis, SemiBin2 and COMEBin, are both based on a deep learning model^12,13^. While, to our knowledge, neither method was trained directly on isolate genomes, it is likely that future methods would be. In such cases, a careful benchmark design that ensures generalization to taxa not seen during training would be necessary.

Overall, MAG-E provides a framework for MAG pipeline evaluation that is flexible, robust, and systematic. Investigators can easily apply MAG-E to other ecosystems, which may have different features (e.g., diversity, abundance distribution), leading to better performance by a different set of tools than those highlighted here. Method developers could use MAG-E to benchmark their tools and detect performance bottleneck for future work. Future applications of MAG-E could include wider benchmarking of different quality control and filtering tools; benchmarking of binning parameters and dereplication performance; and application of MAG-E for evaluation of genome language models.

## Methods

### Metagenomic simulation with MAG-E

MAG-E aims to “mirror” a given metagenomic sample in both strain content and abundance distribution. It requires as input a MAG-E reference database that is built from a collection of MAG and isolate genomes as well as any number of input samples, potentially from a specific ecosystem. In this work, we used the UHGG^1^ v2.0.2 along with a cohort of gut metagenomic samples^29^. MAG-E offers assembly and binning APIs, allowing any user or developer to include and use algorithms of their preference. MAG-E is available with open source from https://github.com/korem-lab/MAG-E, providing instructions on how to construct a reference database, use new assembly, binning, and quality control methods, run analyses, and interpret the results.

### Constructing a MAG-E reference database

A MAG-E reference database is a collection of MAG and isolate genomes that are clustered on two hierarchical levels - a species (95% ANI) and strain (98%) level. To construct it, MAG-E takes as input a collection of genomes preclustered at the species level, for example using dRep^47^ with a 95% ANI threshold, with a representative genome specified for each cluster. MAG-E then further clusters the genomes in each non-singleton species-level cluster to the strain-level using dRep^47^ v3.6.2 “dreplicate”, with the default secondary clustering algorithm and with the ANI cutoff set to 98%. Finally, we use Sylph^25^ v0.9.0 to generate one database of all species-level cluster representatives and another of all genomes in the collection, both built using the Sylph “sketch -c 200” command.

In this work, we used UHGG^1^ is available clustered at the species-level using a 95% ANI. Briefly, Almeida et al.^1^ assigned each genome in UHGG to a random group called a “chunk” that contained 50,000 genomes. Chunks were then independently clustered with dRep using parameters “-pa 0.9 -sa 0.95 -nc 0.30 -cm larger”. From each cluster, a representative was selected, preferring isolates over MAGs, and using a score function accounting for completeness, contamination, and N50 when choosing between isolates (or between MAGs). These representatives were collected across chunks and then clustered together again at 95% ANI. All genomes were then merged into the same cluster as their representative, producing a final species-level clustering and a final species-level representative was selected from each cluster. We downloaded each genome feature format file (.gff) using the ftp links provided in the file genomes-all-metadata.tsv, which we obtained from https://ftp.ebi.ac.uk/pub/databases/metagenomics/mgnify_genomes/human-gut/v2.0.2/. We then extracted the fasta elements from each .gff into a genome-specific fasta file.We used the cluster assignments and species-representatives from the genomes-all-metadata.tsv file as input to MAG-E.

### The MAG-E simulator

For each input sample, MAG-E first constructs a mirror specification, which reflects its metagenomic content using genomes from the MAG-E database, such that the strain structure and species abundance are retained. To construct a specification, we first run the Sylph^25^ “sketch -c 200” command on the input sample, and then run it against the MAG-E species-level Sylph database using Sylph “profile”, providing us with the species-level abundances of the sample. We then run Sylph “query” on the sample’s sketch against the Sylph database of all genomes, providing an adjusted ANI estimate for how well the sample matches each genome in the MAG-E database^25^. For each species with non-zero abundance, we then examine the query results to evaluate whether the sample contains more than one of its strains. We define potential strain genomes as those with an estimated k-mer coverage (Eff_lambda) different than “LOW” (as the ANI estimation can be inaccurately reported in these cases) and with an estimated k-mer-coverage-adjusted ANI (Adjusted_ANI) ≥99.8%. Additionally, if k-mer coverage is not “HIGH”, we also require that the lower bound of the confidence interval on the estimated adjusted ANI ≥99.5%. We group the potential strain genomes by strain cluster, and select one strain from each cluster to add to the specification. To select among potential strain genomes of the same cluster, we prioritize isolates over MAGs, and secondary to this, genomes with higher N50 and ANI. We allow a maximum of three clusters, and, if more clusters are identified, we choose three at random. We then set each of the strains’ abundance to Sylph’s relative abundance estimate for the species multiplied by logNormal(µ=1,σ=2) / S, where S is the sum of the logNormal draws (one draw per strain). If there are no potential strain genomes identified (i.e., adjusted ANI <99.8% for all genomes in the MAG-E database), we select one genome from the species cluster and add it to the specification, prioritizing isolates, then adjusted ANI, and finally N50, and set its abundance to Sylph’s relative abundance estimate.

After we obtain a mirror specification for a sample, we simulate a metagenome for it using InSilicoSeq^26^ v.2.0.1, using the command “generate –draft draft_list –abundance_file abundance_list –n_reads num_reads –model HiSeq” where draft_list is a list of each genome fasta in a specification, abundance_list is the sequence abundance of each genome in a specification, and num_reads is the total number of reads in the real sample.

### The MAG-E manager

The MAG-E manager is the component responsible for running each MAG pipeline, consisting of assembly and binning, as well as for running quality control and binning refinement methods. Three interfaces are exposed to the user: AssemblyTask, BinTask, and QCTask, allowing to extend MAG-E to additional assembly, binning, and refinement / QC algorithms. For assembly, we implemented interfaces and used MEGAHIT^32^ v1.2.9 and MetaSPAdes^33^ v4.2.0. For binning, we implemented and used CONCOCT^10^ v1.10, MaxBin2^9^ v2.2.7, METABAT2^8^ v2.12.1, VAMB^11^ v5.0.4, SemiBin2^13^ v2.2.0, and COMEBin^12^ v1.0.4. For binning refiners, we implemented an interface for DAS Tool^34^ v1.1.7, which, like individual binners, is structured as a BinTask. For quality control and refinement, we implemented and used CheckM2^18^ v1.1.0 and GUNC^38^ v1.0.6. Most binners, including the six we used, require coverage or count information. To generate these, MAG-E maps the reads from each sample against assemblies using Bowtie2^48^ v2.5.4 with default setting in paired end mode, and constructs coordinate-sorted bam files with Samtools^49^ v1.22.1.

MAG-E enables running each binner in one of three modes: (1) “Single-sample” mode, in which only the target sample is used; (2) “Multi-sample” mode, in which that target sample is binned in the context of a dataset with a user defined size (the same “context” dataset is maintained for each sample across the different MAG pipelines evaluated); and (3) “Partial multi-sample” mode, previously used^2^, which is similar to multi-sample mode, except instead of using all samples, we use the D samples closest to the target sample in terms of Mash distance^31^ (by default, D=20).

Following assembly, we construct the ground-truth mapping for each sample. We first construct a BLAST^27^ database using the genomes in the sample’s mirror specification (using the command “makeblastdb -dtype nucl”). We then map each assembly from the sample against its BLAST database using the “blastn” command. We use as ground truth alignments that are highly similar, which we define as percent identity ≥99% and alignment length between 99-101% of the contig length.

### The MAG-E evaluator

MAG-E calculates performance metrics for each ground-truth genome in each sample at the base-pair level. Note that in multi-sample modes, we calculate these only for the target sample. To calculate these metrics, first, for each ground-truth genome and each output bin, we calculate true positives (TP) as the number of ground-truth genome basepairs that are covered by contigs in that bin that map to it (using BLAST described above), and false negatives (FN) as the number of genome basepairs that are not covered. We then select the bin with the highest recall (TP/(TP+FN)) for each ground-truth genome as its representative bin. Once representative bins are assigned, we calculate false positives (FP) as the number of basepairs in contigs in the representative bin that do not map to the ground-truth genome, allowing us to compute precision (TP/(TP+FP)) and F-score (2*recall*precision / (recall + precision)). Note that each genome is assigned exactly one representative bin, but multiple genomes may be assigned to the same bin.

As previously noted^15^, many genomes in realistic samples are not assembled and binned well, resulting in very low recall. To address this, some have advocated for calculating performance metrics with respect to the assembled portion of each genome, which may result in unrealistic performance estimates (discussed in ^35^; e.g., 100% recall for a bin containing a single, 4,000 basepair long contig that was assembled for a particular microbe). This approach is also not applicable to MAG-E, which evaluates entire MAG pipelines including the assembly stage. We address this differently: in **Figs. 2-4**, we only analyze ground-truth genomes that were well recovered (recall 0.7, precision ≥ 0.9) by at least one method; in **Fig. 5**, we only analyze ground-truth genomes with a minimum expected coverage ≥2.5x, which we selected by examining recall vs. expected genome coverage across all ground-truth genomes and across binners (**Fig. S13**), finding a systematic increase in recall for all genomes at around 2.5 regardless of MAG pipeline.

### Metagenomic sample processing

In this work we used gut metagenomic samples from an Israeli cohort^29^ publicly available from ENA, PRJEB11532. We used all 676 samples sequenced on a Illumina HiSeq 2500 platform. Reads were quality filtered with Trimmomatic^50^ v0.39, which, after adapter removal, clipped bases at the start or end of reads with phred ≤25 and dropped reads <50 basepair after quality-based trimming. Reads were then mapped against the GRCh38, CHM13v2, and PhiX genomes with Bowtie2^48^ v2.5.3, and unmapped reads were retained. We then selected samples with ≥5×10^6^ reads for further analysis, leaving 575 samples with a mean±std of 12.43×10^6^±5.98×10^6^ read pairs.

### Benchmarking of the MAG-E simulator to CAMISIM

We constructed simulated metagenomes from the 575 quality-filtered human gut metagenomes using CAMISIM^30^ v1.3-fcc198 and the MAG-E simulator. For CAMISIM, we constructed taxonomic profiles of each real sample using MetaPhlAn^51^ v4.1.0 and ran CAMISIM with community design mode using the “taxonomic-profile-based” command with default arguments, except for setting the thread number to 16 and 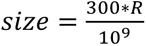 (R - sample read count).

### MAG-E benchmarking

In this work, we evaluated 48 MAG pipelines (two assemblers x six binners + two DAS Tool configurations x six binning modes) on 100 samples we selected at random from the 575 quality-filtered gut metagenomic samples (**Table S13**), resulting in an evaluation of a total of 4,800 individual MAG-generation tasks. All assemblers and binners were run with default settings and according to the original authors’ recommendations for each binning mode. For multi-sample and partial multi-sample modes, we simulated the scenario where each target sample is being binned in the context of a 50-sample dataset by randomly selecting 49 other samples from the 575. Note that while each of the 100 samples was part of a different 50-sample dataset, these datasets were the same across the MAG pipelines.

### Statistical analysis

The evaluation results produced by MAG-E are highly structured: the performance metrics are calculated per ground-truth genome, with many such genomes originating from the same sample, and the same reference genome may be present in multiple samples. Furthermore, different MAG pipelines share steps. While, for visualization purposes, some plots show full data distributions (**Figs. 2a,b, S2a, S3, S5**), our statistical analysis considers the performance over each genome using linear mixed models: for the analysis comparing isolates vs. MAGs (**Fig. 1b**), we used the model “F-score ∼ is_isolate + (1|binner) + (1|binning_mode) + (1|assembler) + (1|sample) + (1|genome)”; for the analysis of MAG pipelines (**Figs. 2, S2**), we used the model “metric ∼ binner * assembler * binning_mode + (1|sample) + (1|genome)”, and the same model is used for **Figs. 3, S6** treating DAS(Nayfach) and DAS(Best) as a binner; for the analysis of contig group recall across binners (**Fig. 4c**), we used the model “recall ∼ contig_group * binner + (1|sample) + (1|genome)”; for the analysis of contig group recall comparing binning modes (**Fig. 4d-f**), we used the model “recall ∼ contig_group * binning mode + (1|sample) + (1|genome)”; and for the analysis of CheckM2 **(Fig. 5a,b,e,f**), we used the model “metric ∼ CheckM2_MIMAG * GUNC_passed * binner * assembler * binning_mode + (1|sample) + (1|genome)”. For calculating R^2^ values in **Fig. 5c,d,g,h** we used the models “recall ∼ CheckM2_completeness + (1|sample) + (1|genome)”, and “contamination ∼ CheckM2_contamination + (1|sample) + (1|genome)”, and reported the marginal R^2^ from the function r2 of the package “performance”^52^. We reported *p* values calculated from two-sided Wald z-test for the relevant contrast in each analysis, adjusted with Tukey Honestly Significant Difference. Analyses were performed using the lme4^53^ v1.1-38, emmeans^54^ v2.0.1, and performance^52^ v0.16.0 packages in R^55^ v4.5.2.

### Contig-level analyses

To analyze the effect of contig properties on binning, we analyzed each contig from the set of recoverable genomes. First, we computed the cosine distance between the normalized tetra-nucleotide composition of each contig and that of its ground-truth genome. We then also calculated the Euclidean distance between each contig’s coverage vector and the mean coverage vector from all contigs assigned to the same ground-truth genome as the contig, with both vectors calculated across the dataset. If a contig was shared among multiple genomes in the recoverable set, its properties were considered for each ground-truth genome it mapped to. We then discretized these distances into percentiles at the basepair level. To do this, we ordered the contigs from all samples by distance and calculated the cumulative sum of their lengths. A contig c was assigned to the p-th percentile (p from 0-99), if the cumulative sum up to the contig s_c_ plus half its length l_c_ was p*bases < s_c_ + l_c_ / 2 ≤ (p+1)*bases, where bases is the sum of basepairs in all contigs of the recoverable genomes / 100. We then calculated (FN) TP for the contigs in each percentile in each sample as the number of basepairs from contigs in that percentile that were (in)correctly binned in the sample, i.e., were assigned to the bin that (did not) represents the ground-truth genome of the contig.

To stratify contigs as belonging to the accessory / variable genome (**Fig. 4c-f**), we ran the “end-to-end” command of geNOMAD^36^ v1.11.2 on each genome in the recoverable genome set, and recorded high-confidence provirus annotation (virus score ≥0.9). We then classified each contig as prophage if, in one of the genomes it mapped to, it was entirely contained within a prophage, contained an entire prophage, or if at least 25% of a prophage was covered by the contig. We also classified contigs as shared if they mapped (via BLAST, described above) to two or more recoverable genomes. To evaluate different binners, we then calculated the recall of each group (all contigs, prophage, or shared) in each recoverable genome in a sample.

## Data availability

The gut metagenomic dataset^56^ is available from ENA, accession PRJEB11532. The MAG-E database for UHGG will be made available on Zenodo.

## Code availability

MAG-E is available on GitHub at https://github.com/korem-lab/MAG-E.

## Acknowledgments

We thank members of the Korem group for useful discussions. We thank Jingqiu Liao for early discussions about this project. The study was supported by the Program for Mathematical Genomics at Columbia University and R01AI183668 (T.K.).

## Author contributions

I.C., G.Q. and T.K. conceived the study. I.C. and T.K. designed the study, designed analyses, interpreted the results, and wrote the manuscript with input from all authors. I.C. wrote MAG-E and performed all analyses, with assistance from J.M., G.Q., Y.S., and A.B.K. T.K. supervised the study.

## Supplementary figures

**Supplementary Figure 1.**
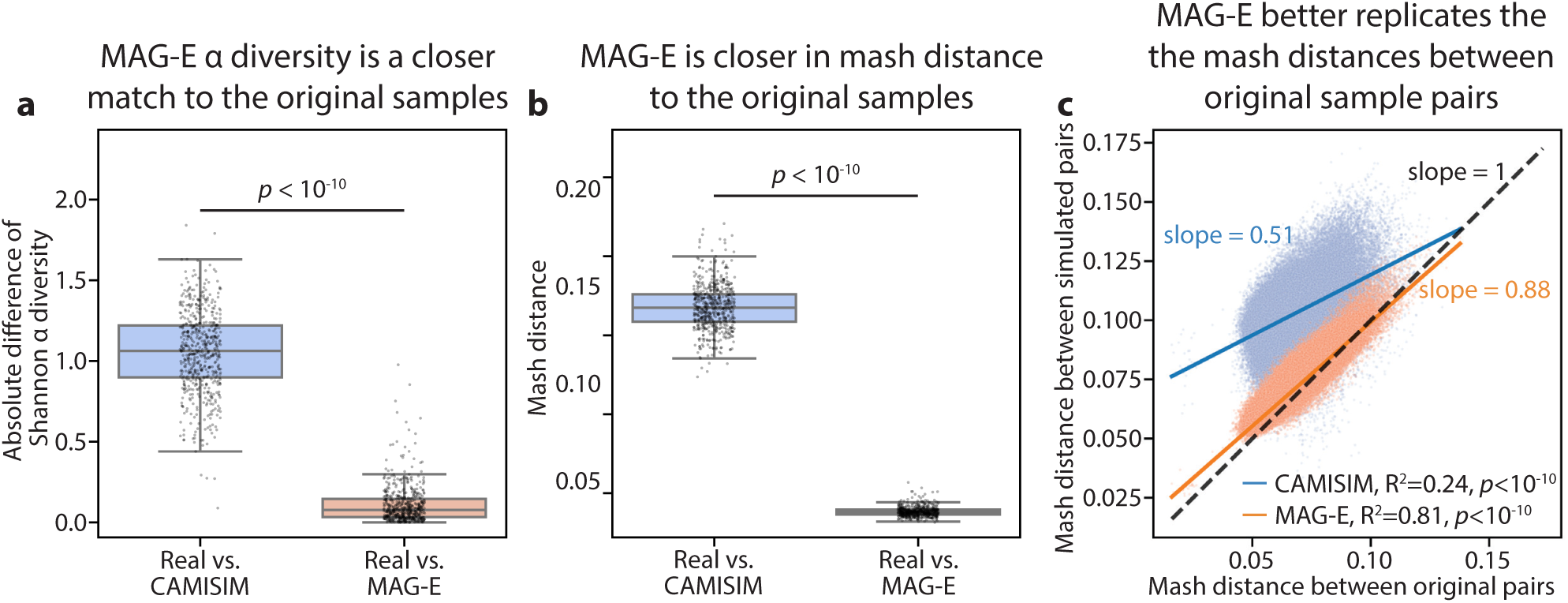
**a,** Difference in Shannon α diversity between simulated and original metagenomes for 575 human gut metagenomes, comparing between CAMISIM and MAG-E. **b-c,** Same as Fig. 1e**,f**, respectively, using Mash distance instead of β diversity. Box, IQR; line, median; whiskers, 1.5xIQR; *p*, Wilcoxon signed-rank test (a,b) or Student’s t-test (c).

**Supplementary Figure 2.**
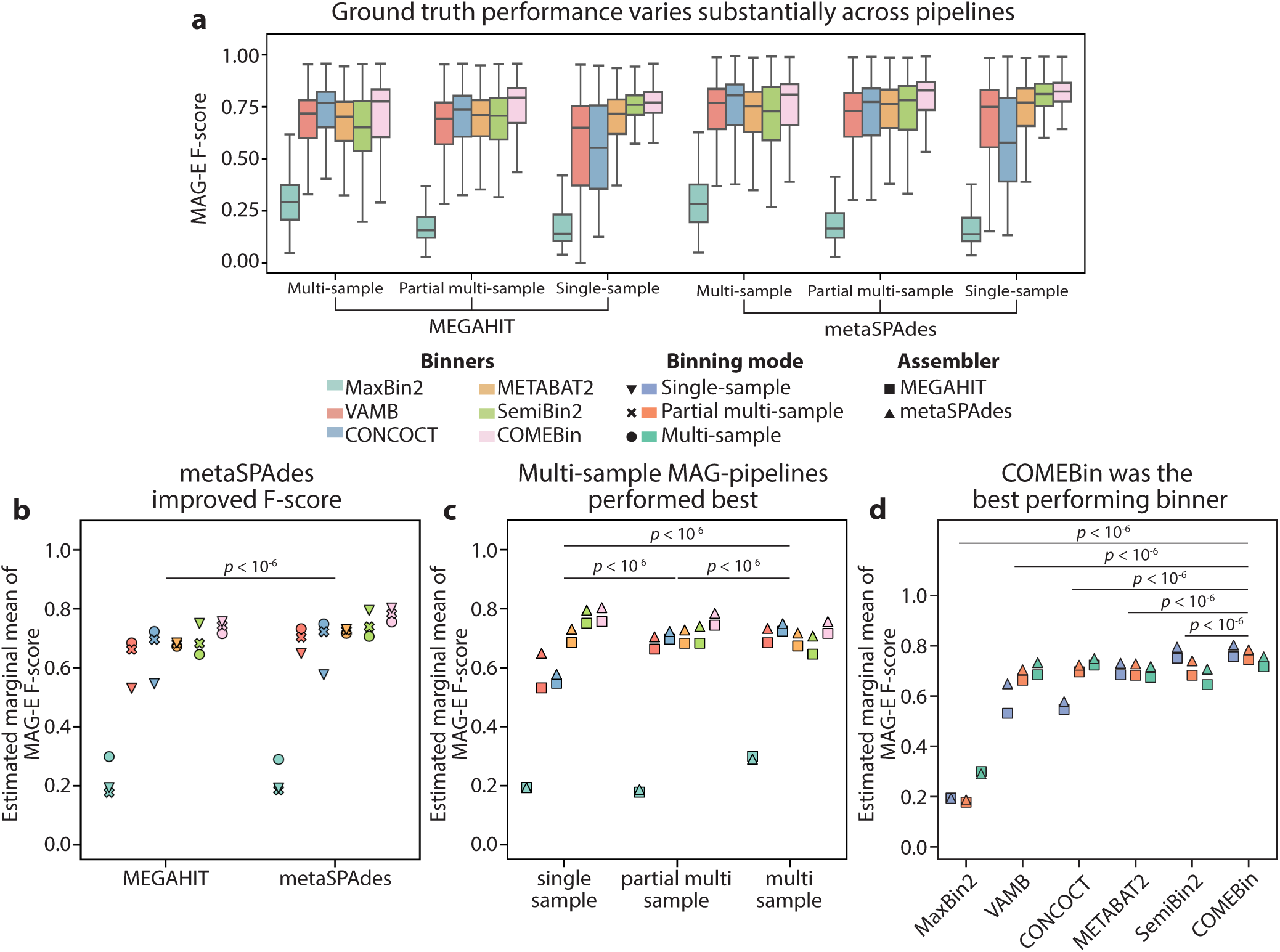
The MAG pipeline of metaSPAdes followed by multi-sample COMEBin outperforms alternatives based on F-scores. **a-d**, Same as **Fig. 2a,c,e,g**, respectively, showing F-score instead of recall.

**Supplementary Figure 3.**
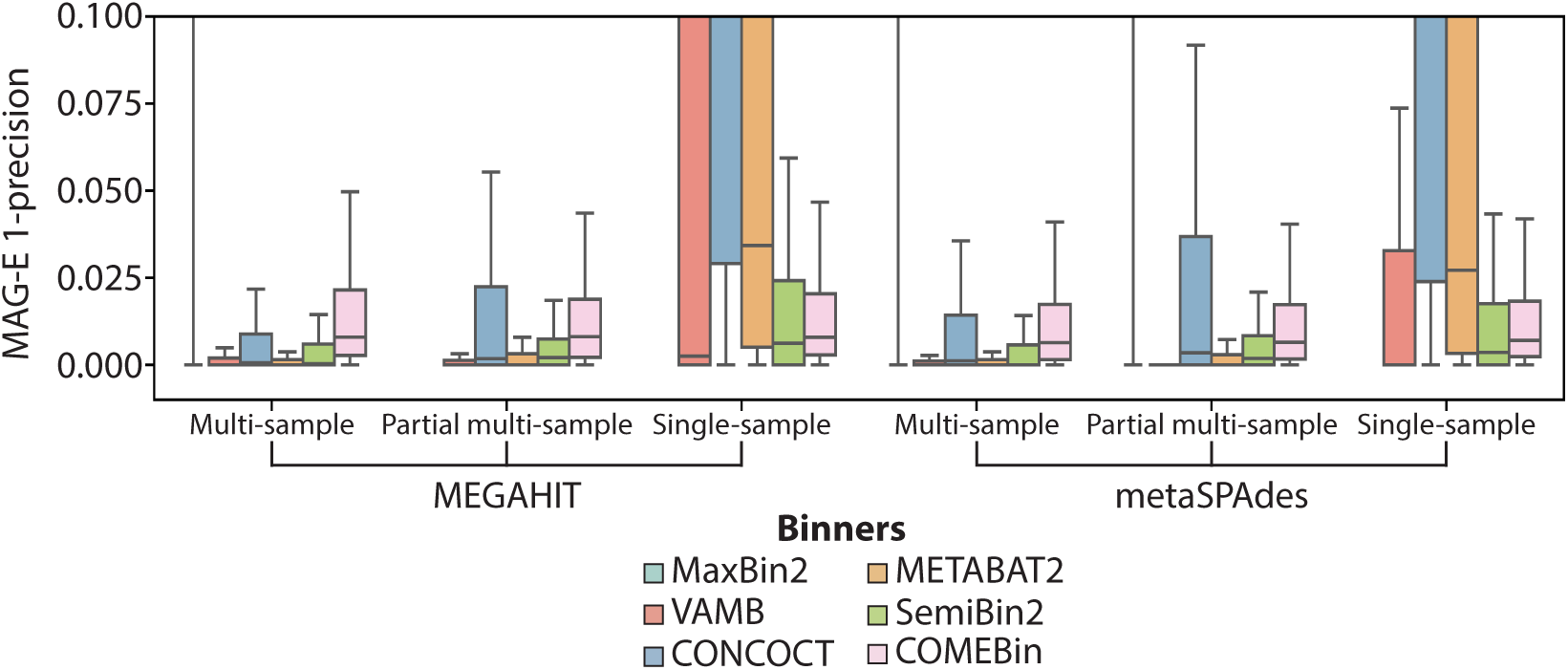
1-Precision of MAG pipelines over the [0, 0.1] range. Box, IQR; line, median; whiskers, 1.5xIQR.

**Supplementary Figure 4.**
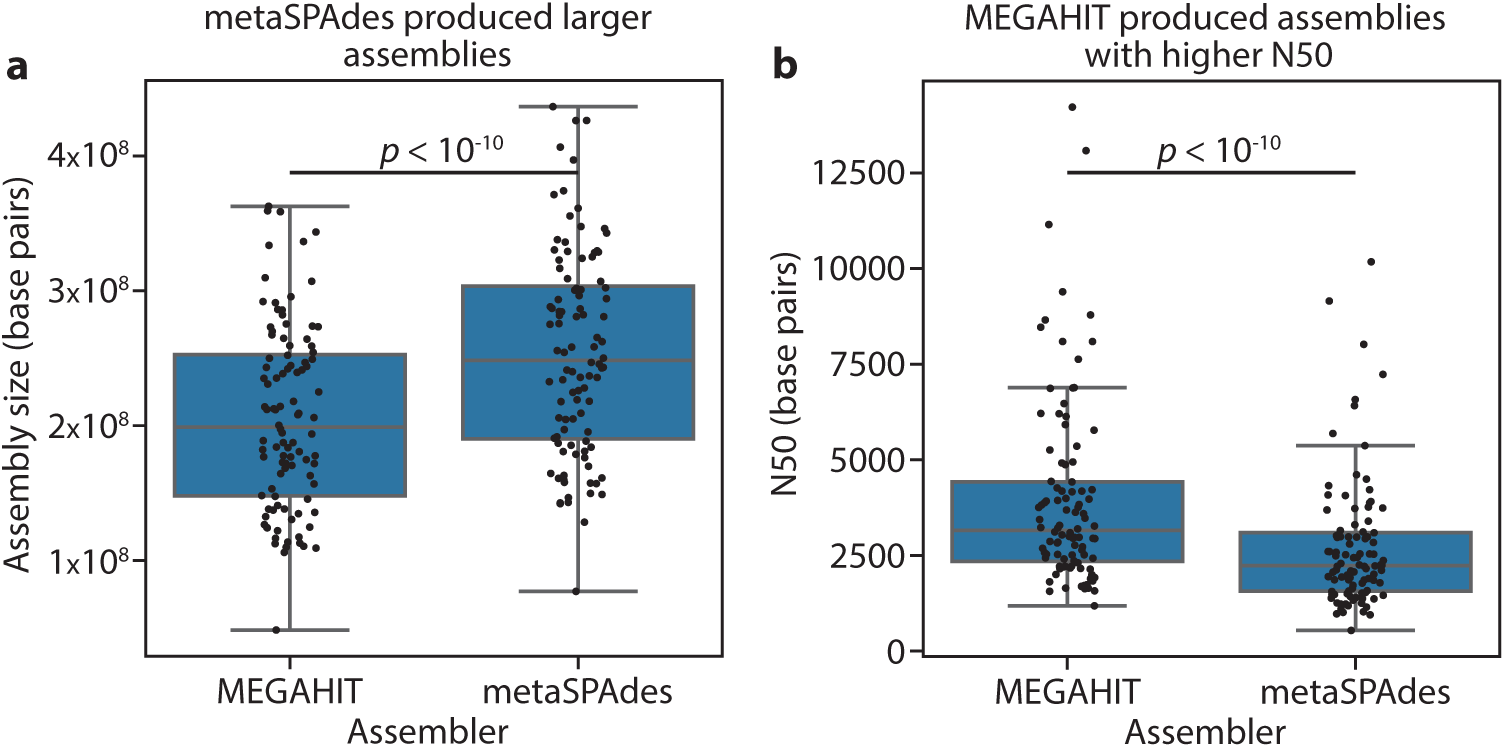
Assembly statistics for metaSPAdes vs MEGAHIT on the 100 samples evaluated with MAG-E. N50 (**a**) and assembly size (**b**; measured as the total number of basepairs) of each MEGAHIT and metaSPAdes assembly. Boxplot, IQR; line, median; whiskers, 1.5xIQR; *p*, Wilcoxon signed-rank test.

**Supplementary Figure 5.**
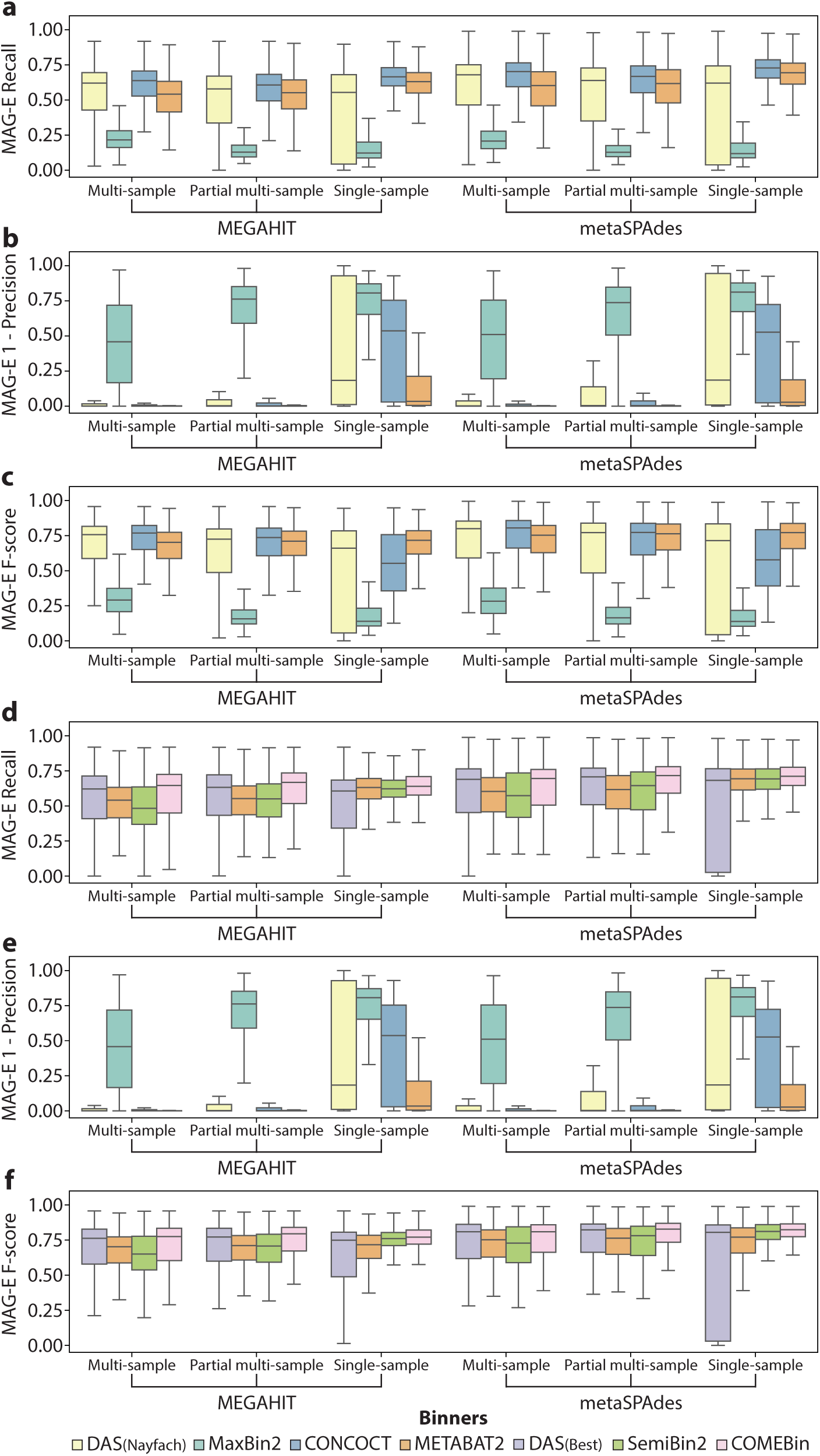
MAG-E recall, 1-precision and F_score for DAS Tool and associated binners. **a-c,** Boxplots showing MAG-E recall (a), 1-Precision (b) and F-score (c) of the recoverable genomes across all samples for DAS(Nayfach) and the binners it combines. **d-f**, same as (a-c) respectively, but for DAS(Best) and the binners it combines. Box, IQR; line, median; whiskers, 1.5xIQR.

**Supplementary Figure 6.**
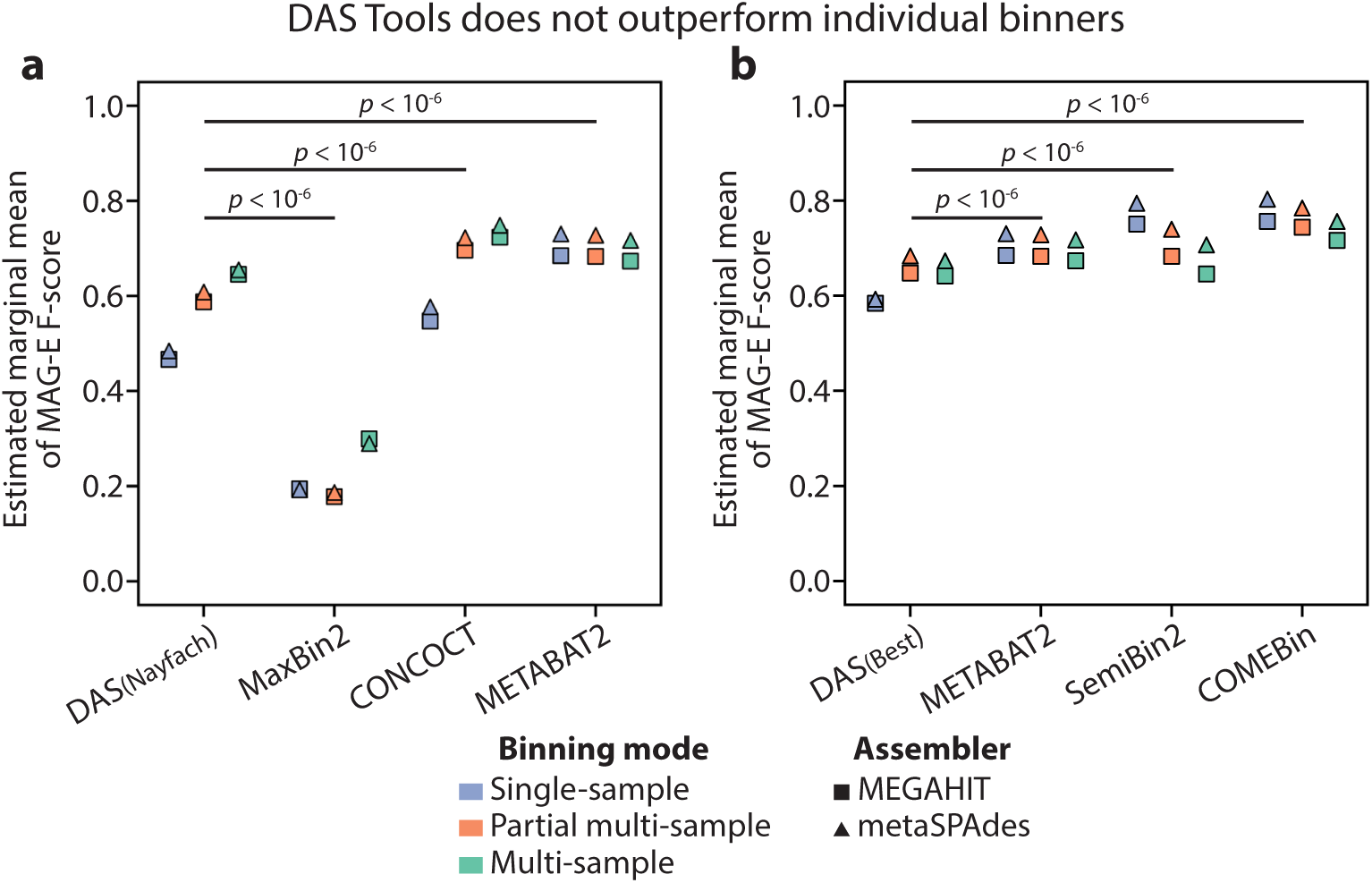
Applying DAS tools does not improve performance over individual binners. **a,b,** Same as Fig. 3a**,c**, respectively, showing F-score instead of recall. Two-sided Wald z-test, *p*<10^−6^, Tukey HSD adjusted.

**Supplementary Figure 7.**
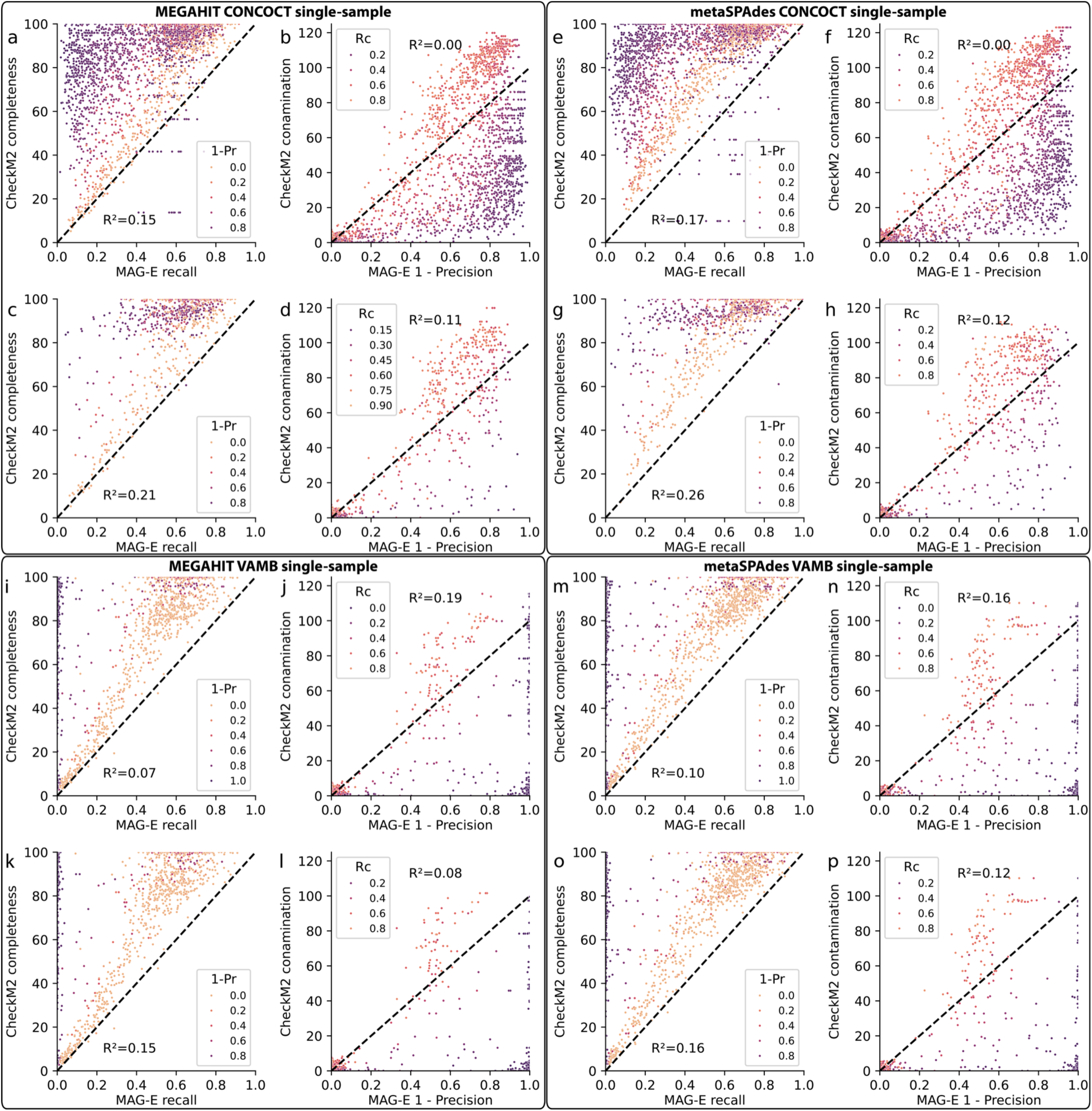
Evaluation of CheckM2 completeness and contamination versus ground-truth, per-genome MAG-E evaluation for CONCOCT and VAMB in single-sample mode. **a-p,** Same as Fig. 5c**,d****,g,h** for CONCOCT after MEGAHIT (a-d) or metaSPAdes (e-h) assembly, or VAMB following MEGAHIT (i-l) or metaSPAdes (m-p) assembly. Rc, recall; 1-Pr, 1-precision; black dashed line, y=100x, indicating perfect correspondence between checkM2 and MAG-E; R^2^, marginal explained variance from a linear mixed model of ground truth from checkM2 predictions (**Methods**).

**Supplementary Figure 8.**
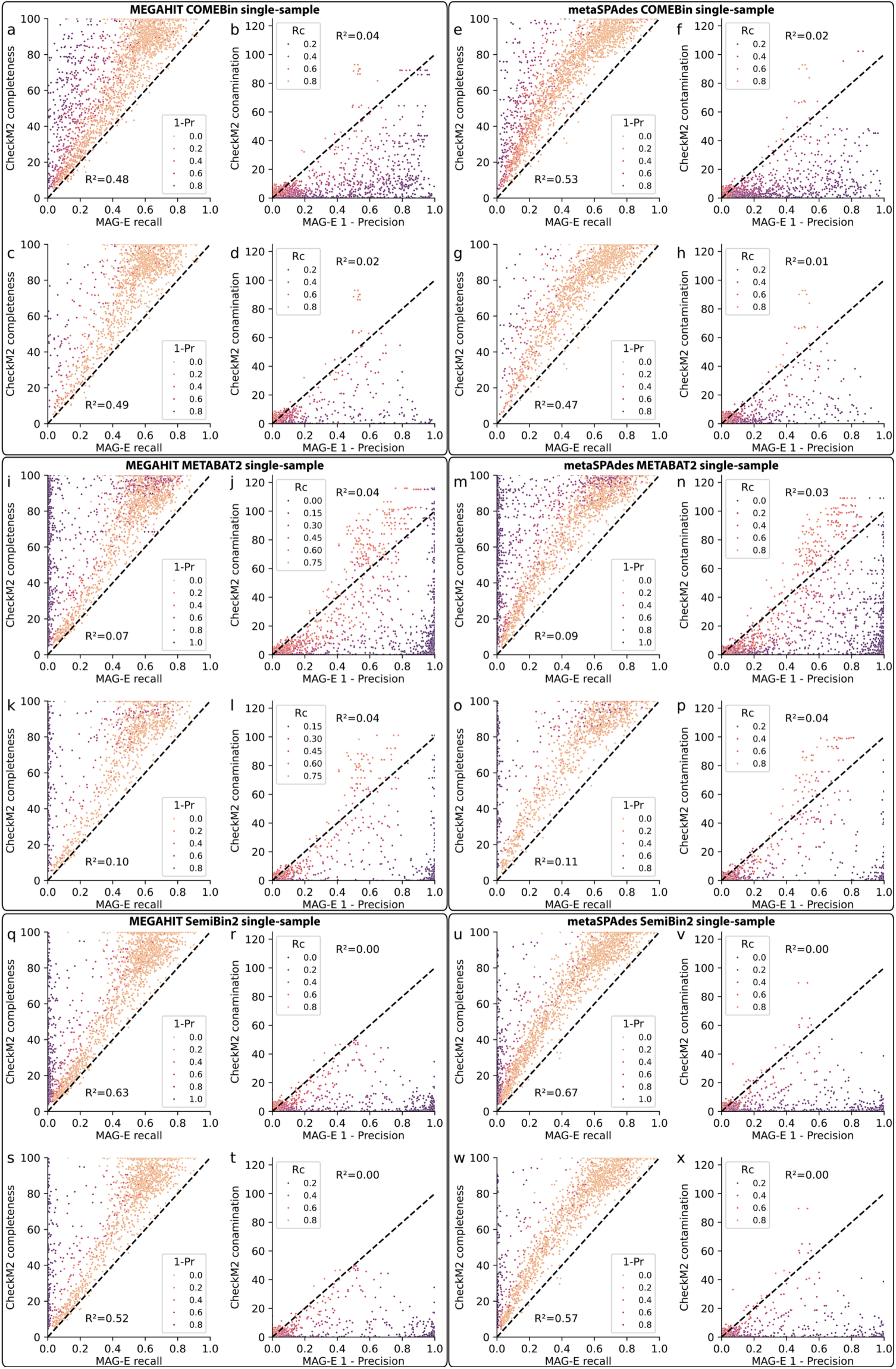
Evaluation of CheckM2 completeness and contamination versus ground-truth, per-genome MAG-E evaluation for COMEBin, METABAT2, and SemiBin2 in single-sample mode. **a-x,** Same as Fig. 5c**,d****,g,h** for COMEBin after MEGAHIT (a-d) or metaSPAdes (e-h) assembly, METABAT2 binning after MEGAHIT (i-l) or metaSPAdes (m-p) assembly, or SemiBin2 following MEGAHIT (q-t) or metaSPAdes (u-x) assembly. Rc, recall; 1-Pr, 1-precision; black dashed line, y=100x, indicating perfect correspondence between checkM2 and MAG-E; R^2^, marginal explained variance from a linear mixed model of ground truth from checkM2 predictions (**Methods**).

**Supplementary Figure 9.**
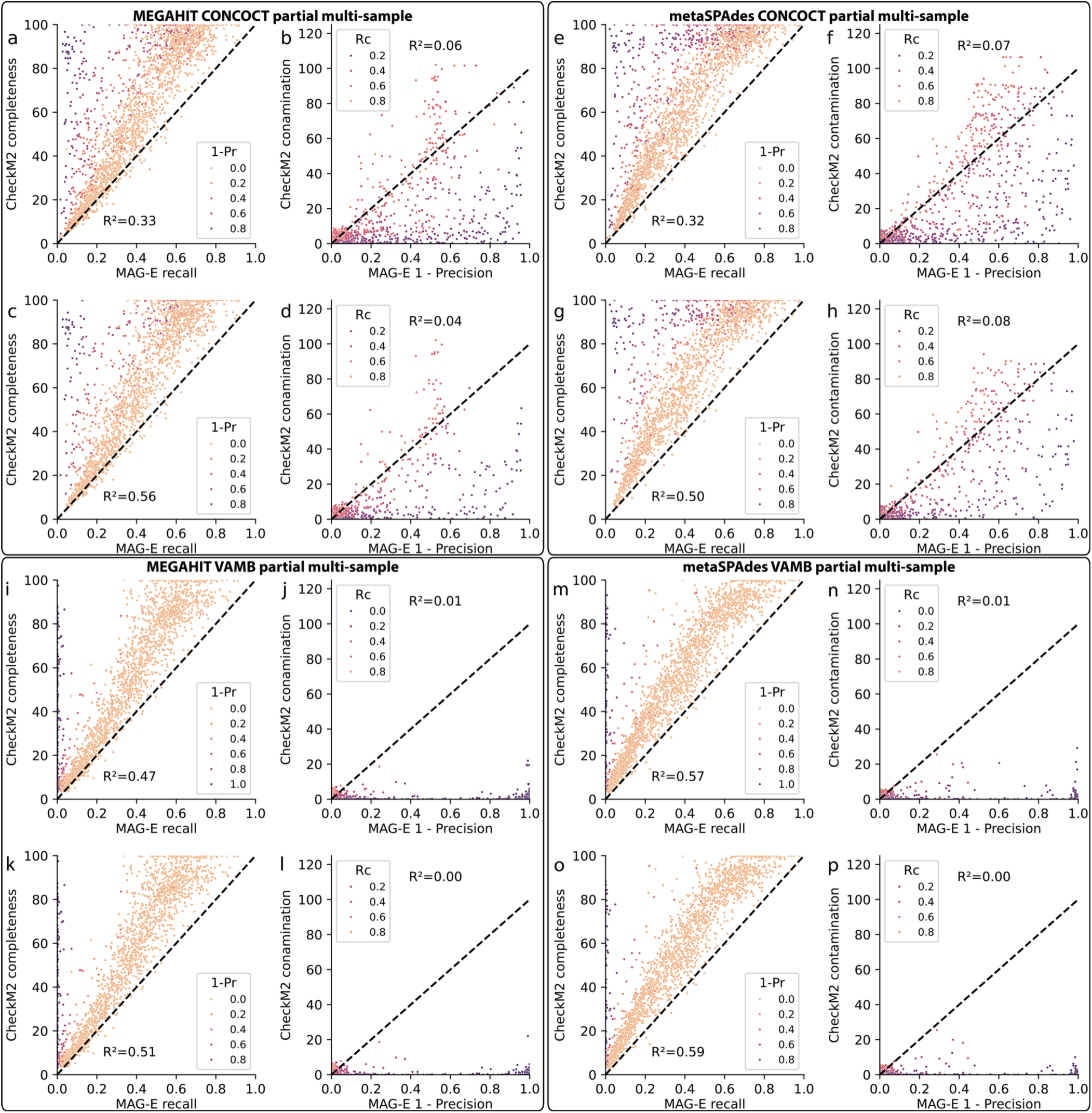
Evaluation of CheckM2 completeness and contamination versus ground-truth, per-genome MAG-E evaluation for CONCOCT and VAMB in partial multi-sample mode. Same as **Fig. S7** for partial multi-sample mode.

**Supplementary Figure 10.**
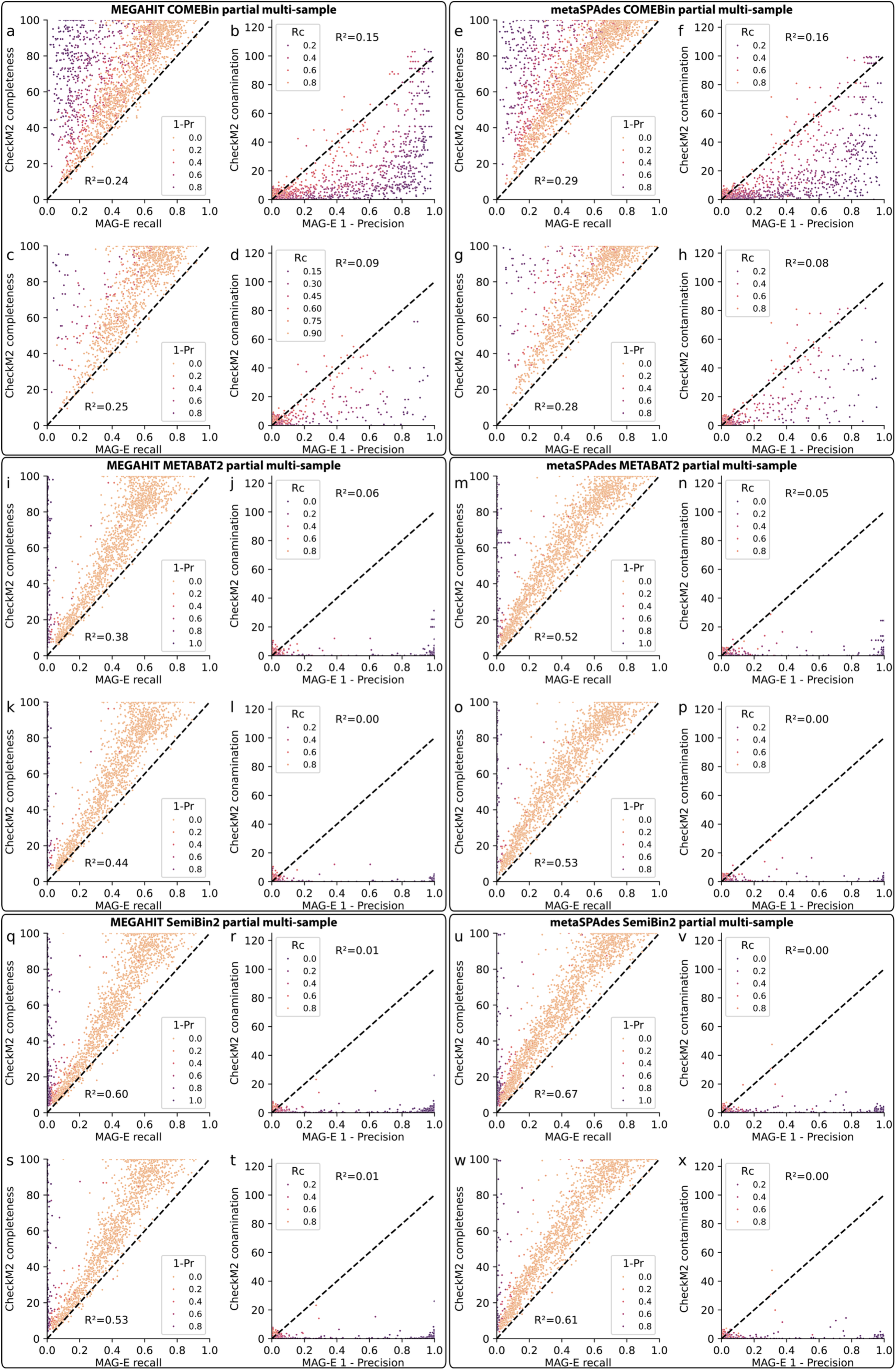
Evaluation of CheckM2 completeness and contamination versus ground-truth, per-genome MAG-E evaluation for COMEBin, METABAT2, and SemiBin2 in partial multi-sample mode. Same as **Fig. S8** for partial multi-sample mode.

**Supplementary Figure 11.**
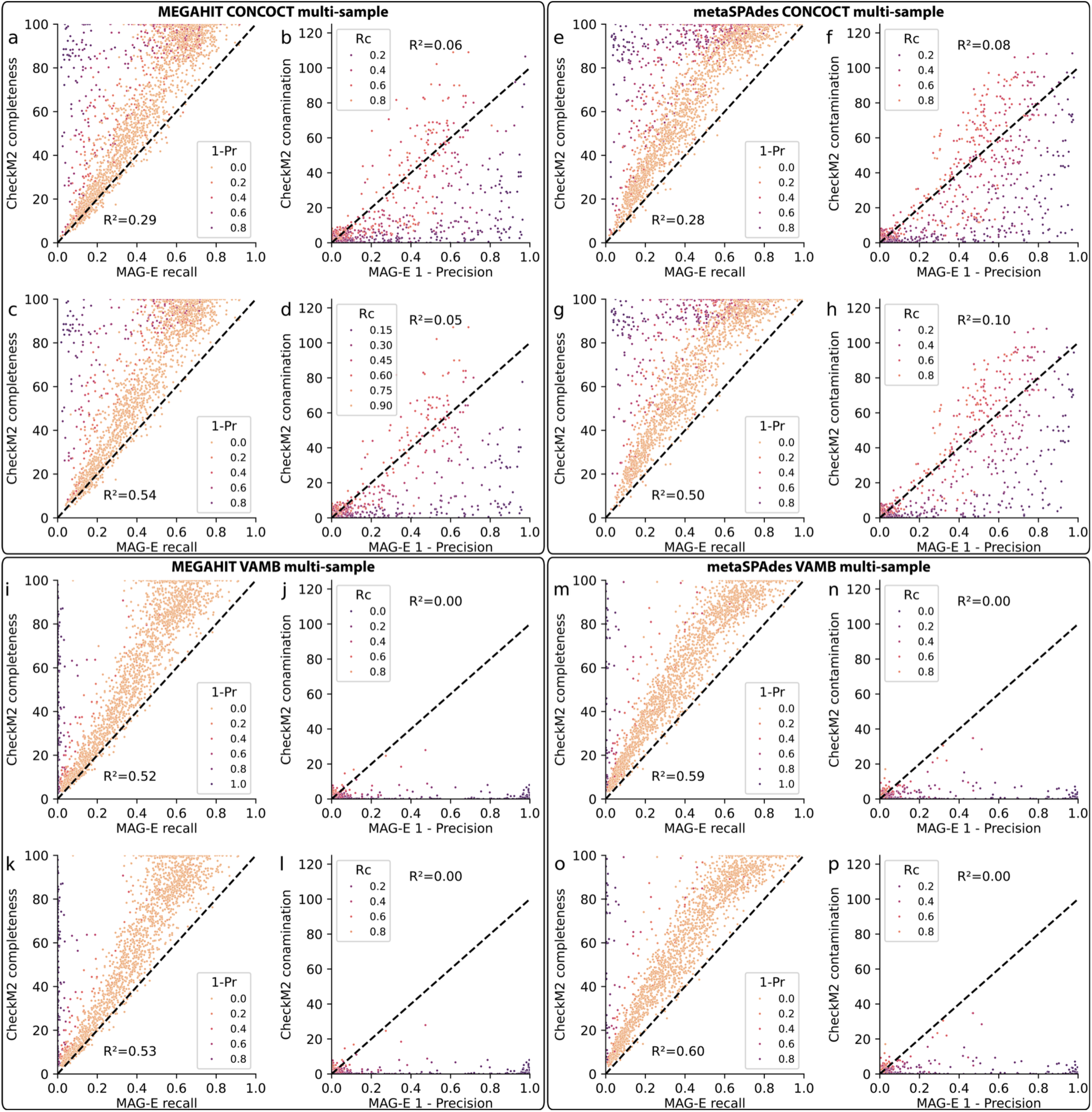
Evaluation of CheckM2 completeness and contamination versus ground-truth, per-genome MAG-E evaluation for CONCOCT and VAMB in multi-sample mode. Same as **Fig. S7** for multi-sample mode.

**Supplementary Figure 12.**
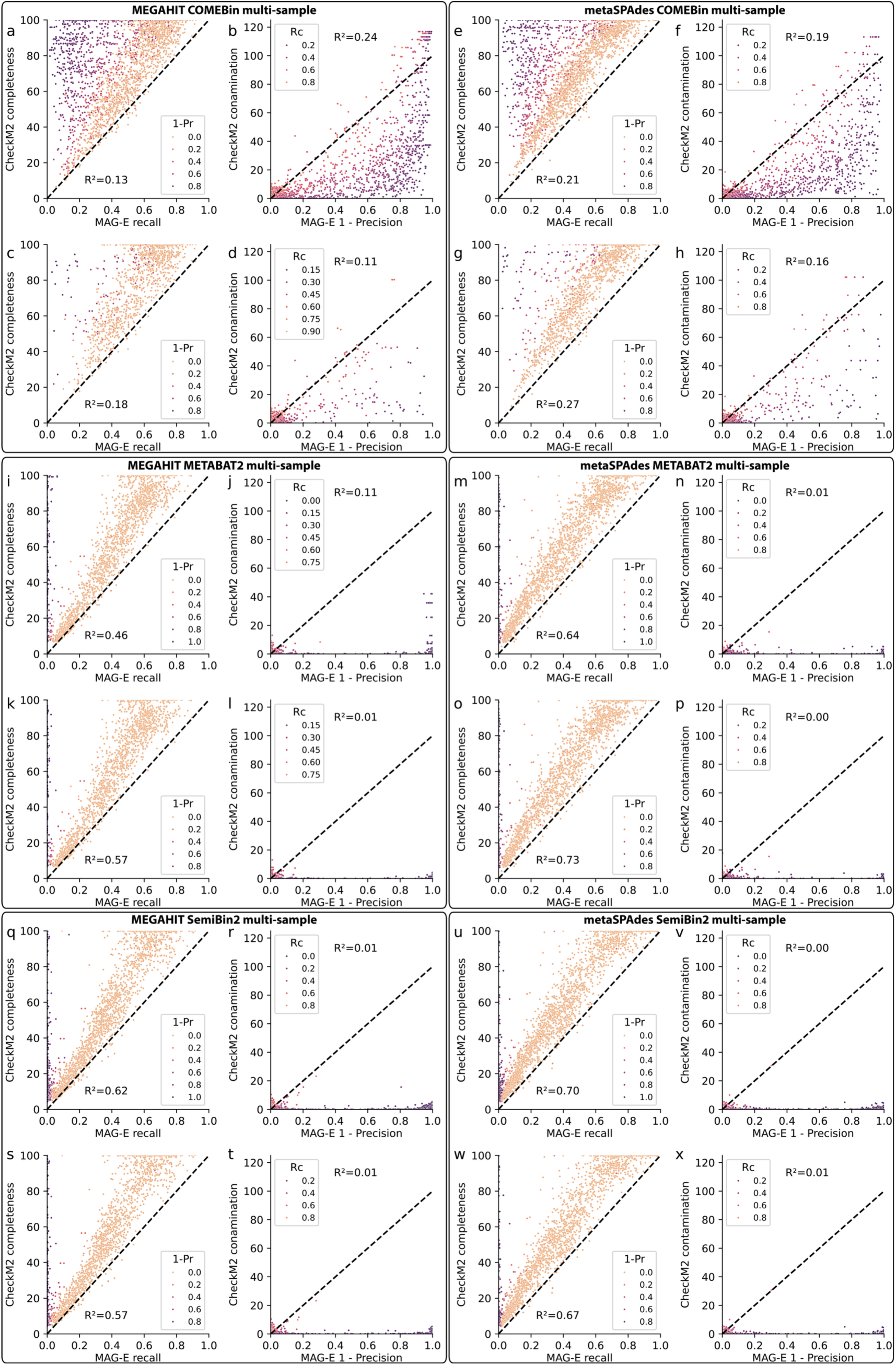
Evaluation of CheckM2 completeness and contamination versus ground-truth, per-genome MAG-E evaluation for COMEBin, METABAT2, and SemiBin2 in multi-sample mode. Same as **Fig. S8** for multi-sample mode.

**Supplementary Figure 13.**
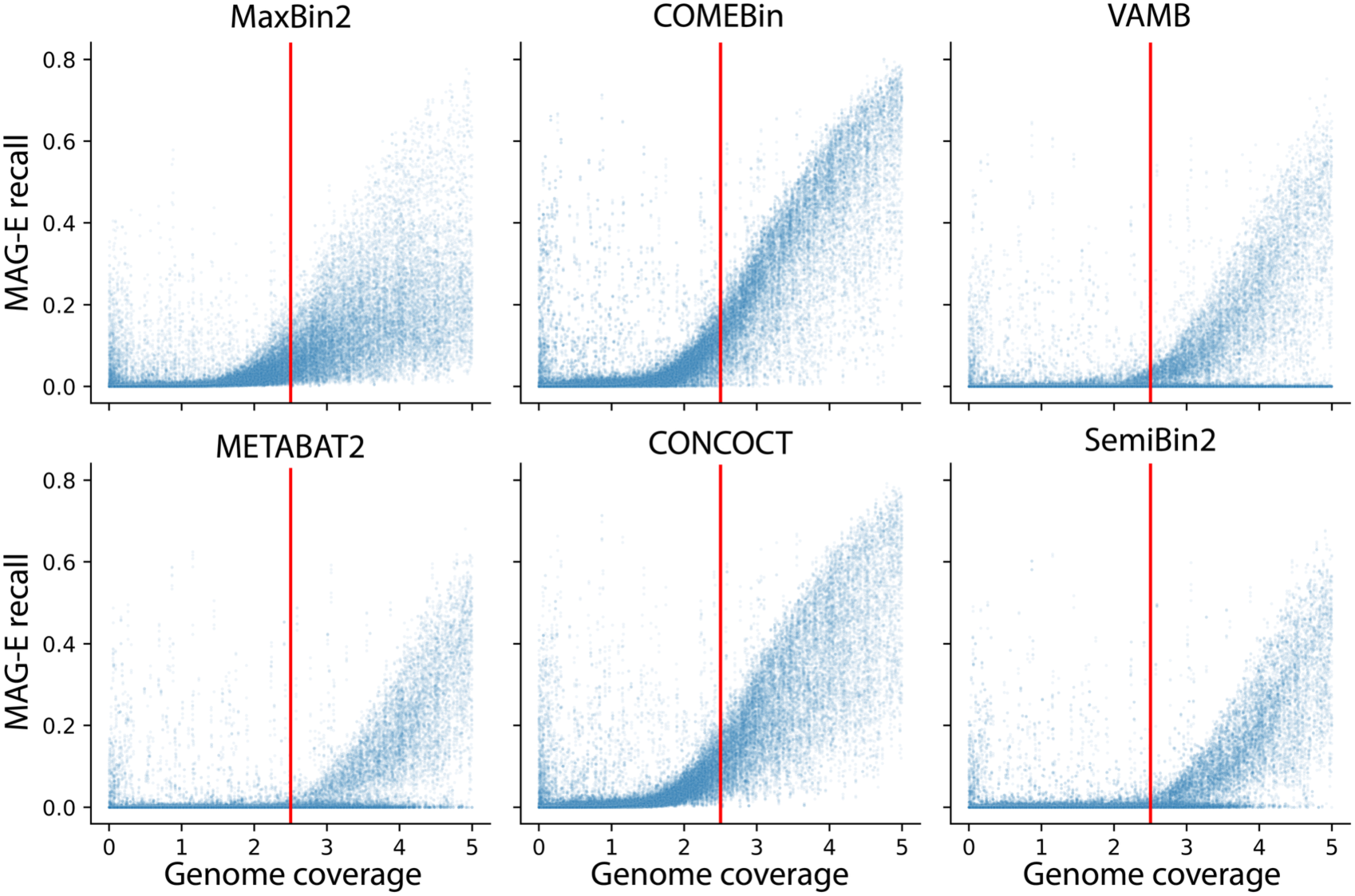
Recall (y-axis) vs. expected coverage (x-axis) for all ground-truth genomes across the different MAG-pipelines, stratified by each binner, demonstrating a substantial increase after 2.5x coverage (red lines).

## Supplementary tables

**Supplementary Table 1.** Summary of past binning benchmark studies. Listed for each study is: whether the metagenomic datasets considered of simulated or real sequencing reads; whether the exact genomic sequence that the MAGs derive from was known, i.e., whether a ground truth was used; dataset complexity, describing the microbiome source or number of elements in the dataset; number of samples in the dataset; whether the analysis examined the binning of contigs with particular properties; evaluated tools used to quantify MAG quality; and compared the performance of different short-read metagenomic assemblers, binning modes, and binners in combination, rather than just the performance of binning.

| Study (dataset) | Simulated or real | Ground truth | Dataset complexity | Dataset size (samples) | Contig-level | Quality filtering | MAG pipeline |
| --- | --- | --- | --- | --- | --- | --- | --- |
| Sczyrba et al., 2017<sup>21</sup> (high complexity) | Simulated | Yes | 596 genomes, 478 circular elements | 1 | No | No | No |
| Sczyrba et al., 2017<sup>21</sup> (medium complexity) | Simulated | Yes | 132 genomes, 100 circular elements | 1 | No | No | No |
| Sczyrba et al., 2017<sup>21</sup> (low complexity) | Simulated | Yes | 40 genomes, 20 circular elements | 1 | No | No | No |
| Meyer et al., 2022<sup>22</sup> (marine) | Simulated | Yes | 77 genomes, 200 circular elements | 10 | No | No | No |
| Meyer et al., 2022<sup>22</sup> (plant) | Simulated | Yes | 496 genomes, 398 circular elements | 21 | No | No | No |
| Meyer et al., 2022<sup>22</sup> (strain madness) | Simulated | Yes | 408 genomes | 100 | No | No | No |
| Maguire et al., 2020<sup>24</sup> | Simulated | Yes | 30 genomes, 65 plasmids | 1 | Yes | Yes | No |
| Nelson et al., 2020<sup>19</sup> (mock) | Real | Yes | 20 genome mock community | 1 | Yes | No | No |
| Nelson et al., 2020<sup>19</sup> (tara oceans) | Real | Partially | Marine microbiome | 1293 | Yes | No | No |
| Mattock et al., 2023<sup>17</sup> | Real | No | Cow rumen | 42 | Yes | Yes | No |
| Han et al., 2025<sup>23</sup> | Real | No | Marine, cheese, human, sludge | 5 | No | No | No |
| Yue et al., 2020<sup>44</sup> (chicken gut) | Real | No | Chicken gut | 4 | No | No | No |
| Yue et al., 2020<sup>44</sup> (simulated) | Simulated | Yes | CAMI 1 high, medium, low complexity | 3 | No | No | No |

**Supplementary Table 2**. Recall, precision, and F-score of all isolate genomes and MAGs for each of the 36 pipelines, presented in **Fig. 1b**.

**Supplementary Table 3**. MAG-E recall, precision, and F-score of recoverable genomes across 100 samples for each of the 48 MAG pipelines, presented in **Figs. 2a,b, S2a, S3,** and **S5a-f**.

**Supplementary Table 4.** Marginal mean MAG-E recall estimated by linear mixed models (**Methods**) for each of the 48 MAG pipelines, presented in **Figs. 2c,e,g** and **3a,c**.

**Supplementary Table 5.** Marginal mean MAG-E precision estimated by linear mixed models (**Methods**) for each of the 48 MAG pipelines, presented in **Figs. 2d,f,h** and **3b,d**.

**Supplementary Table 6.** Marginal mean MAG-E F-score estimated by linear mixed models (**Methods**) for each of the 48 MAG pipelines, presented in **Figs. S2b-d and S6a,b.**

**Supplementary Table 7.** Contig-level recall of the all contigs, prophage elements, and shared elements of the recoverable genomes across 100 samples for METABAT2, SemiBin2 and COMEBin run in multi-sample mode following assembly by metaSPAdes, presented in **Fig. 4c**.

**Supplementary Table 8.** Contig-level recall of all contigs, prophage elements, and shared elements of the recoverable genomes across 100 samples for METABAT2 run in single-sample, partial multi-sample and multi-sample mode, presented in **Fig. 4d**.

**Supplementary Table 9.** Contig-level recall of all contigs, prophage elements, and shared elements of the recoverable genomes across 100 samples for SemiBin2 run in single-sample, partial multi-sample and multi-sample mode, presented in **Fig. 4e**.

**Supplementary Table 10.** Contig-level recall of all contigs, prophage elements, and shared elements of the recoverable genomes across 100 samples for COMEBin run in single-sample, partial multi-sample and multi-sample mode, presented in **Fig. 4f**.

**Supplementary Table 11.** Recall, precision, and F-score of the recoverable genomes from each of 30 pipelines classified into MIMAG categories by CheckM2, with and without GUNC filtering, presented in **Fig. 5a,b,e,f.**

**Supplementary Table 12.** Recall, precision, F-score, CheckM2 completeness and CheckM2 contamination for all genomes ≥ 2.5x coverage that were successfully binned, presented in **Figs. 5c,d,g,h, S7-12**.

**Supplementary Table 13**. List of 100 metagenomic samples used for evaluation of MAG pipelines in this work.

